# Improved segmentation of the intracranial and ventricular volumes in populations with cerebrovascular lesions and atrophy using 3D CNNs

**DOI:** 10.1101/2020.03.23.000844

**Authors:** Emmanuel E. Ntiri, Melissa F. Holmes, Parisa M. Forooshani, Joel Ramirez, Fuqiang Gao, Miracle Ozzoude, Sabrina Adamo, Christopher J.M. Scott, Dar Dowlatshahi, Jane M. Lawrence-Dewar, Donna Kwan, Anthony E. Lang, Sean Symons, Robert Bartha, Stephen Strother, Jean-Claude Tardif, Mario Masellis, Richard H. Swartz, Alan Moody, Sandra E. Black, Maged Goubran

## Abstract

Successful segmentation of the total intracranial vault (ICV) and ventricles is of critical importance when studying neurodegeneration through neuroimaging. We present iCVMapper and VentMapper, robust algorithms that use a convolutional neural network (CNN) to segment the ICV and ventricles from both single and multi-contrast MRI data. Our models were trained on a large dataset from two multi-site studies (N=528 subjects for ICV, N=501 for ventricular segmentation) consisting of older adults with varying degrees of cerebrovascular lesions and atrophy, which pose significant challenges for most segmentation approaches. The models were tested on 238 participants, including subjects with vascular cognitive impairment and high white matter hyperintensity burden. Two of the three test sets came from studies not used in the training dataset. We assessed our algorithms relative to four state-of-the-art ICV extraction methods (MONSTR, BET, Deep Extraction, FreeSurfer), as well as a ventricular segmentation tool (FreeSurfer). Our multi-contrast models outperformed other methods across all evaluation metrics, with average Dice coefficients of 0.98 and 0.94 for ICV and ventricular segmentation respectively. Both models were also the most time efficient, segmenting the structures in orders of magnitude faster than some of the other available methods. Our networks showed an increased accuracy with the use of a conditional random field (CRF) as a post-processing step. We further validated both segmentation models, highlighting their robustness to images with lower resolution and signal-to-noise ratio, compared to tested techniques. The pipeline and models are available at: https://icvmapp3r.readthedocs.io and https://ventmapp3r.readthedocs.io to enable further investigation of the roles of ICV and ventricles in relation to normal aging and neurodegeneration in large multi-site studies.

## Introduction

Variant patterns of cerebral atrophy are observed in patients suffering from various neurodegenerative diseases, such as cerebrovascular disease (CVD), Parkinson’s disease (PD), and Alzheimer’s disease (AD) (Staffaroni et al. 2017; Burton et al. 2004). Patients with ischemic stroke have cerebral atrophy in regions connected to the brain lesion (Kraemer, Schormann, and Hagemann 2004). Other overt vascular brain lesions such as ischemic infarcts, as well as covert lesions of presumed vascular origin such as white matter hyperintensities (WMH), exacerbate brain tissue atrophy and increase the risk of cognitive decline (Stebbins et al. 2008; E. E. Smith et al. 2015; Swartz et al. 2008). Brain atrophy can be observed as *in vivo* longitudinal changes on structural MRI, including decreased total brain volume and increased ventricular size (Aribisala et al. 2012). Differential rates and patterns of structural brain atrophy are key biomarkers for cognitive decline and vascular cognitive impairment in both research and clinical settings (Boccardi 2003; Carmichael et al. 2007; Nestor et al. 2008; Chou et al. 2010; Madsen et al. 2015). Identifying changes in structural biomarkers such as total brain and ventricular volumes can serve as a means of tracking disease progression and risk of conversion to dementia.

Intracranial volume (ICV) is defined as the total volume of grey matter, white matter, and cerebrospinal fluid (CSF) (Huo et al. 2017). ICV is commonly used to correct for head size when evaluating volumetric or morphometric properties of brain tissue (Mathalon et al. 1993; S. M. Smith et al. 2001), and is assumed to be unaffected by neurodegeneration (Jenkins et al. 2000). Brain volume normalized by ICV has been found to differentiate between dementia groups and cognitively normal older individuals (Wolf et al. 2004; Bigler and Tate 2001). It is also associated with cognitive reserve after adjusting for the presence of pathology (van Loenhoud et al. 2018). Skull stripping, the segmentation step that extracts the ICV by differentiating between its representative voxels and surrounding tissue, is often the first step in a vast majority of neuroimaging pipelines. Hence, its accuracy is critical for a wide range of downstream structural and functional analyses.

The ventricular system consists of the lateral ventricles, the third and fourth ventricle, and foramen that connect the three. Ventricular size serves as a biomarker for cognitive decline with age (Apostolova et al. 2012). A large body of literature has investigated correlates between the ventricular system and different neurodegenerative diseases. A study conducted by Yang et al. demonstrated that morphometric properties of the ventricles accurately predicted conversion to mild cognitive impairment (MCI) and AD (Yang, Tan, and Qiu 2012). In addition, different trajectories of volumetric expansion have been found between cognitively normal individuals and patients with frontotemporal dementia (FTD) and AD (Driscoll et al. 2009; Thompson et al. 2004; Whitwell et al. 2015).

Cerebrovascular lesions including white matter hyperintensities (WMH) and ischemic strokes are concomitant in both cognitively normal adults and those suffering from neurodegenerative diseases (van Dijk et al. 2002). For example, patients post-stroke have a substantially increased risk of dementia (Ivan et al. 2004). High WMH burden is associated with lower processing speed and impaired cognitive performance (Dong et al. 2015), and patients with both increased WMH and ventricular volumes perform worse in several cognitive domains including episodic memory, semantic memory, and processing speed (Dong et al. 2015). The presence of these vascular lesions, along with brain atrophy, in neurodegenerative populations necessitates the need for accurate algorithms that segment key brain structures in the presence of cerebrovascular injury.

Automated methods of segmenting the ICV and ventricles have been developed for use in neuroimaging pipelines, but these methods may not produce optimal segmentations in patients with vascular brain lesions or atrophy (Fischl et al. 2002; Stephen M. Smith 2002; Chou et al. 2009; Kleesiek et al. 2016; Roy et al. 2017). Despite the importance placed on these imaging biomarkers in neuroscience, segmentation algorithms are commonly developed using young, healthy individuals due to the limited number of large studies with aging control populations. In individuals with significant atrophy, delineating the boundary of brain tissue can be difficult (Fennema-Notestine et al. 2006). Many established segmentation methods also do not account for or produce suboptimal results in the presence of infarction or WMH, which are concomitant in populations with vascular brain injury (Breteler et al. 1994; Aribisala et al. 2012). WMH localizes around the ventricular horn and can be difficult to differentiate from ventricular cerebrospinal fluid (CSF) on T1-weighted scans. Although some methods may produce accurate segmentations in the presence of vascular lesions or atrophy, they are often computationally inefficient and require extensive parameter tuning or manual intervention to deal with anatomical heterogeneity.

As newer studies strive to collect larger amounts of neuroimaging data from multiple cohorts across multiple sites, there is an increased need to develop robust and fast whole brain and ventricular segmentation methods (Ross et al. 2015). Early segmentation methods relied on intensity or template-based approaches (Carmichael et al. 2007; Chou et al. 2010; Madsen et al. 2015). In recent years, machine learning based algorithms, and in particular convolutional neural networks (CNNs), have gained traction for brain segmentation methods with promising results (Kamnitsas et al. 2017; Roy et al. 2017; Kayalibay, Jensen, and van der Smagt 2017; Kleesiek et al. 2016). A CNN is a type of neural network that can handle complex computer vision problems because it uses multiple layers at once much like the visual system of the brain (Lecun et al. 1998) and has been shown to be quite accurate in tasks such as classification and image segmentation (Krizhevsky, Sutskever, and Hinton 2017). In a supervised learning setting, by using a dataset of ground-truth labels (often manually traced), CNNs extract the underlying features in an image that results in the optimal segmentation. During training the network optimizes a loss function to generate the output with the lowest (or highest) loss compared to the ground-truth. A commonly used CNN architecture in biomedical applications is a 3D U-Net structure which combines features extracted at lower and higher levels for a more accurate segmentation (Tajbakhsh et al. 2016; Ronneberger, Fischer, and Brox 2015).

In this work, we introduced and evaluated CNN-based algorithms to segment the ICV and ventricles in the presence of vascular lesions and atrophy using a large dataset (N=528 subjects for ICV, N=501 for ventricular segmentation) from three multi-site studies; encompassing clinically complex populations with a large variability in brain anatomy due to aging, stroke, and neurodegeneration. Our algorithms were trained on semi-automated segmentations meticulously edited by trained experts. We trained these algorithms on different combinations of input sequences to determine the sequence effect or weighting on segmentation. We compared our methods against established state-of-the-art (SOTA) techniques to highlight their accuracy and efficiency. The initial segmentation results were further compared with post-processed outputs generated by a conditional random field (CRF). Finally, to further validate the performance of our models, they were assessed against simulated challenges in clinical datasets, namely noise and reduced resolution, referred to here as “clinical adversarial attacks”. Our models are publicly available, and we developed an easy-to-use pipeline with a graphical interface for making them accessible to users without programming knowledge.

## Materials and Methods

### Participants

To train the ICV and ventricle segmentation models, a total of 539 participants were recruited from 2 multicenter studies: 214 subjects with cerebrovascular disease +/- vascular cognitive impairment (CVD +/- VCI) or Parkinson’s disease (PD) (55-86 years, 72% male) through the Ontario Neurodegenerative Disease Research Initiative (ONDRI), and 325 individuals with non-surgical carotid stenosis (47-91, 63% male) through the Canadian Atherosclerosis Imaging Network (CAIN) (ClinicalTrials.gov: NCT01440296). Of the participants recruited, 528 were used to train the network for iCVMapper, and 501 were used to train VentMapper after exclusions through manual quality control of the data and ground truth segmentations. Ground truth segmentations for the ICV and ventricles were generated using the SABRE and Lesion Explorer semi-automated pipeline that generate intensity-based segmentations which are then manually edited by expert raters trained by a neuroradiologist with an intraclass correlation of ≥ 0.9 (Ramirez et al. 2011; Ramirez et al. 2014). Sequences used for generation of the ground truth data included 3D T1-weighted, T2-weighted fluid attenuated inversion recovery (FLAIR) and an interleaved T2-weighted and proton density (T2/PD) sequence to accurately delineate the brain parenchyma and ventricular CSF. Participant demographics, diagnosis, Montreal Cognitive Assessment (MoCA) scores, and volumes for the ground truth segmentations and vascular lesions are summarized in **Table 1**.

**Table 1.** Participants study demographics, clinical diagnosis, MoCA scores, ICV and ventricle volumes, as well as WMH and stroke volumes in the training and test datasets. Data is presented as mean ± standard deviation unless otherwise specified.

|  | **Training Dataset**<br>**n = 539 \*** |  | **Test Dataset**<br>**n = 132** |  |  |
| --- | --- | --- | --- | --- | --- |
|  | **CAIN**<br>**n=325** | **ONDRI**<br>**n= 214** | **MITNEC**<br>**n=50** | **ONDRI**<br>**n=82** | **SDS**<br>**n=106** |

| *Demographics* |  |  |  |  |  |
| --- | --- | --- | --- | --- | --- |
| Diagnostic Group | CN | PD, CVD | MCI, AD, CVD | PD | VCI |
| Age (years) | 70.4 (7.9) | 69.0 (7.1) | 76.8 (8.2) | 67.7 (6.3) | 73.8 (8.0) |
| Sex (n, % female) | 117 (36.8) | 60 (28.0) | 23 (46.0) | 19 (23.2) | 48 (45.3) |
| MOCA total score | -- | 26.8 (2.6) | 22.3 (4.8) | 26.2 (2.4) | -- |
| *Neuroimaging Metrics* |  |  |  |  |  |
| ICV (cc) | 1246.4 (128.8) | 1250.6 (134.8) | 1223.9 (146.4) | 1313.8 (134.9) | 1217.7 (122.7) |
| vCSF (cc) | 38.4 (20.1) | 41.2 (22.6) | 55.8 (22.9) | 36.4 (17.1) | 56.9 (26.8) |
| WMH (cc) | 9.0 (10.2) | 8.8 (11.7) | 37.1 (23.7) | 4.9 (5.1) | 18.9 (20.3) |
| Stroke (cc) | 9.7 (14.4)<sup>☆</sup> | 11.7 (21.3)<sup>◇</sup> | 0.4 (0.4)<sup>†</sup> | 0.2 (0.2)<sup>+</sup> | 47.3 (42.0)<sup>*</sup> |

\* 528 used for ICV and 501 for ventricular segmentation after quality control
MoCA=Montreal Cognitive Assessment; vCSF= ventricular cerebrospinal fluid; WMH=white matter hyperintensities; CN=cognitively normal; PD=Parkinson’s Disease; VCI=vascular cognitive impairment; MCI=mild cognitive impairment; AD=Alzheimer’s disease. <sup>☆</sup> available in 57/325 participants; <sup>◇</sup> available in 90/214 participants; <sup>†</sup> available in 5/50 participants; <sup>+</sup> available in 2/82 participants; <sup>\*</sup> available in 38/106 participants.

The models were tested on two datasets from different multi-site studies (**Table 1**): 1) 82 subjects from the ONDRI PD cohort, and 2) 50 subjects with severe WMH burden (greater than Fazekas 2) from the Medical Imaging Trials NEtwork of Canada (MITNEC) Project C6 (ClinicalTrials.gov: NCT02330510), a separate study not part of those used for training, and 3) 106 subjects with VCI characterized by atrophy and vascular lesions from the Sunnybrook Dementia Study (SDS), another study not part of the training set. The acquisition parameters for the sequences used from the four studies can be found in the **Supplementary Materials**. The T1-only model was used for the SDS study since a FLAIR sequence was not available in the study’s scanning protocol.

### Model architecture

Our networks are based on the U-net architecture (Ronneberger, Fischer, and Brox 2015; Çiçek et al. 2016), which consists of contracting and expanding pathways, with pooling and upsampling operators. Features that are passed at each “descending” level are also passed to levels on the opposite end through skip connections, such that information from downsampling levels is utilized in higher levels.

The architecture of the network (**Figure 1**) was based on the original 2D U-net (Çiçek et al. 2016; Ronneberger, Fischer, and Brox 2015), with some modifications in our 3D implementation. Similar to our prior work (Goubran et al. 2019), residual blocks were added to each encoding layer. Residual blocks resolve the gradient degradation problem that occurs with deeper networks with an increasing number of layers. Here, one of our contributions was the incorporation of dilated convolutions. Dilated convolutions help to enlarge the field of view (FOV) of convolutional filters without losing resolution or coverage (Yu and Koltun, 2015). An equally weighted formulation of the Dice coefficient was used as the loss function to mitigate the class imbalance issue (the majority of image voxels do not represent the structure of interest). Instance normalization was used instead of batch normalization (Ioffe and Szegedy 2015) to avoid instability of batch normalization due to the stochasticity generated by small batch sizes. The network had a depth of 5 layers and 16 initial filters.

**Figure 1.**
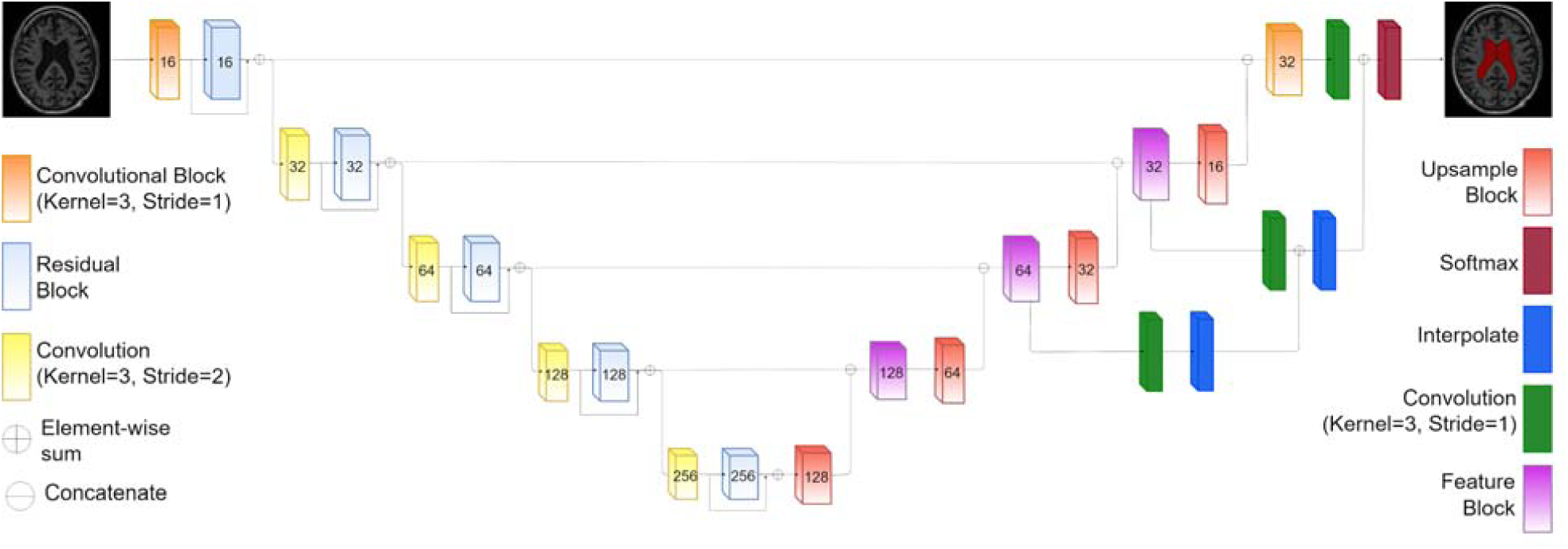
Proposed architecture for the 3D-Unet convolutional neural network with residual blocks and dilated convolutions.

Each contracting step consisted of a convolutional layer and a residual block (He et al. 2016). At each residual block, the input is split into two paths. Dilated convolutions (with a dilation rate of 2) were applied in the first path, while an identity map was applied to the input data in the second path. Element-wise addition was then applied to the results of the first and second paths, combining the outputs of the two. The first path consisted of two dilated convolutional layers with kernels sizes 7×7×7 with a dropout layer with a rate of 0.3 between the two.

At each ascending layer before the final layer, the upsampling block from the previous level is concatenated with the corresponding features on the contracting level. Upsampling modules consisted of an upsampling layer of size 3×3×3, followed by a convolutional layer. The output from the upsampling module was concatenated with the summation output from the respective later on the contracting side, before being passed to a feature block. The feature block consisted of two convolutional layers (one dilated convolution) with a stride of 1×1×1, and kernels of size 7×7×7 and 1×1×1, respectively. The number of filters was halved at each expanding step. At the last layer, the concatenated feature map is passed to a softmax function to generate a probability map for the region of interest (ROI).

### Loss Function

A modification of the Dice coefficient (Dice 1945) was used as a loss function, as initially proposed by Milletari et al (Milletari, Navab, and Ahmadi 2016). The Dice coefficient is a measure of similarity, determined by a calculation of the overlap between two binary images. Given a predicted truth binary volume, *P* and a ground truth binary volume *G*, the Dice coefficient is defined as:

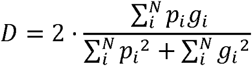

When differentiated with respect to *P*_*j*_, for the back-propagation pass, we get:

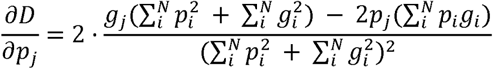

### Data preprocessing, augmentation and model training

The preprocessing techniques used in this study are described in our prior work (Goubran et al. 2019). Briefly, all images used for training were bias-field corrected using *N4* (Tustison et al. 2010). *C3d* was then used to standardize the intensities of the images within a neighborhood of 50×50×50 voxels (Yushkevich et al. 2006). A total of three augmentations per scan were performed on each of the subjects, including flipping on the Left-Right axis, as well as a rotation on the Left-Right axis by 15 and -15 degrees. No deformable augmentations were used. All models were trained for 500 epochs. Early stopping was set to 50 epochs where validation loss did not improve to avoid overfitting. The Adam optimizer (Kingma and Ba 2014) was used with an initial learning rate of 5×10^−3^, and a learning rate drop of 0.5. We have selected these training hyperparameters based on validation experiments and previous literature.

Two networks were trained with different inputs for each segmentation task: a multi-contrast network that relies on T1-weighted, T2-weighted and FLAIR sequences as inputs, and a T1-based network. The networks were trained on images that were downsampled to a size of 128×128×128, and were trained on the whole image, as opposed to patches. Each network was trained with 5-fold cross validation. 10% of the training set was used for validation. The networks were trained on a GeForce GTX1080 Ti graphics card with 11G of memory and a Pascal architecture (NVIDIA, Santa Clara, CA).

### Evaluation of clinical datasets

Our ICV model performance was compared against four other segmentation tools: 1) the Brain Extraction Tool (BET), a widely used segmentation tool that generates a deformable surface around assumed brain tissue (Stephen M. Smith 2002), 2) Multi-cONtrast brain STRipping method (MONSTR) (Roy et al. 2017), a multi-atlas based approach that utilizes multiple sequences and patch-based data from multiple atlases to generate a brain mask, 3) a CNN-based technique (Deep Extraction) that uses an 8-layer fully convolutional CNN using any combination of four sequences to extract the brain from MRI images (Kleesiek et al. 2016), and 4) FreeSurfer (stable version 6.0.0) which uses prior probabilities and atlas registration to calculate the likelihood that an anatomical structure occurs at a given location in atlas space (Fischl et al. 2002; Fischl, Sereno, and Dale 1999). To conduct a fair comparison while accounting for the possibility of confounding the results, our statistical analysis was performed across models with the same number of inputs (i.e. comparing single-contrast models together and multi-contrast together). All components of the ventricular system (i.e. the left and right lateral ventricles, the left and right inferior lateral ventricles, the third and fourth ventricles) were combined and used for comparison. Our ventricular model performance was compared against FreeSurfer’s segmentation, as all the open-source methods that were researched only segmented the lateral ventricles. The default parameters for each segmentation method were used during testing.

Volume and shape-based metrics were used to evaluate the segmentation performance of the segmentation methods including the Pearson correlation coefficient, the Dice correlation coefficient, the Jaccard similarity coefficient, the Hausdorff distance, and the absolute volume difference. Pearson’s correlation coefficient (Pearson and Galton 1895) was used as a measure of correlation between the volumes from each automatic segmentation prediction P, and volumes from ground-truth manually segmented data G. The Pearson correlation coefficients were then transformed to Fisher’s z-transform values. The Jaccard coefficient (Jaccard 1912) is a measure of the similarity between two datasets, calculated by dividing the intersection of the two sets by their union. Given a predicted binary mask, *P*and a binary ground truth volume *G*, the Jaccard coefficient is defined as:

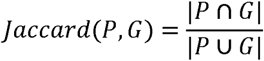

The Hausdorff distance measures how similar two objects occupying the same space are. Given two sets of points representing objects occupying the same space, *A* and *B*, where *x ∈ A* and *y ∈ B*, the Hausdorff distance from *A* to *B* is defined as the largest value in a derived set of closest distances between all points.

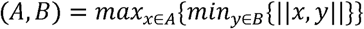

In the above function,| | *x,y* | | is the Euclidean distance between points *x* and *y*. Because the Hausdorff distance between the two sets relative to *A* is not equal to the distance relative to *B*, the bidirectional Hausdorff distance is equal to the maximum value between the two directions. A smaller distance is indicative of a greater degree of similarity between the segmentation and the manual tracing.

The absolute volume difference (AVD) between the volumes of both ground truth and predicted images was also computed. An AVD of 0 signifies that the ground truth and the segmentation have the same number of voxels, though it is not indicative of a perfect segmentation.

The non-parametric Mann-Whitney U test was employed with α-level of 0.05 to assess the improvement between our models and all tested methods on these evaluation metrics. Multi-contrast and T1-based models were compared separately.

### Post-processing segmentation improvement

We employed a fully connected conditional random field (CRF) to further refine the segmentation predictions. A CRF works by maximizing label agreement between similar elements of a set based on contextual features. Consider an image, *X*, and *Y*, a set of random variables with a domain of a finite set of labels *L* = {*L* _1_,*L*_2_,…, *L* _*K*_}, where *X* and *Y* are the same size, *N*. A CRF (X, Y) is defined as a classifier that aims to accurately define the probability P(Y|X), for all neighboring voxels in Y while minimizing an energy function. CRF potentials incorporate smoothness kernels that maximize label agreement between similar pixels, hence enabling more precise segmentation.

Krähenbühl and Koltun (Krähenbühl and Koltun 2011) utilized a fully connected CRF, in which pairwise potentials are found across all pixel pairs in the image, as opposed to solely neighboring pixels. The Gibbs energy of a fully connected pairwise CRF model is defined as:

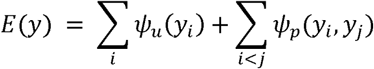

Where *i* and *j* are pixels. The unary potential *ψ*_*u*_(*y*_i_) is calculated independently for each pixel by a classifier given image features including shape, texture, location, and color descriptors. The pairwise potential, *ψ*_*p*_(*y*_i_,*y*_*j*_) takes the form:

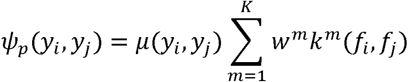

Where *μ*(*y*_i_,*y*_*j*_) is a label compatibility function defined by the Potts model,*k*^*m*^ is a Gaussian kernel, *f*_*i*_ and *f*_*j*_ are feature vectors for pixels *i* and *j*. The CRF used in this study was implemented by Kamnitas et al. (Kamnitsas et al. 2017, 2015), which built on the work by Krahenbuhl and Koltun (Krähenbühl and Koltun 2011) by implementing a 3D version of the CRF, extending the segmentation to medical images.

### Clinical adversarial attacks

In order to validate the robustness of our models on data with lower resolution or quality, we generated adversarial cases to further test the models’ performance. These cases included: downsampling of image resolution (to simulate typically short clinical scans performed on low field strength magnets) and introduction of noise (to simulate data with lower signal-to-noise ‘SNR’ ratio). Input images were downsampled in two ways: a) by a factor of 2 across all dimensions or b) by a factor of 2 in the x and y dimensions, and a factor of 4 in the z-dimension. Noise was introduced using a gamma distribution and standard deviations of σ={0.1, 0.3, 0.5}. Other SOTA methods were also compared to our models on the most challenging adversarial cases, those downsampled by a factor of 2, 2 and 4 across the x, y and z dimensions respectively, as well as those with induced noise of σ=0.5.

## Results

In this section, we first compared the segmentation performance of the models when trained on different combinations of the input sequences (T1, T2 and FLAIR), in order to determine the sequence importance for the segmentation task. We then compared the performance of our models to SOTA methods using Pearson’s correlation, Dice, Jaccard coefficients and Hausdorff distance, as well as processing time. We then applied a CRF to refine the segmentation predictions and assess whether results could be further improved. The models were finally tested against generated adversarial cases to evaluate their robustness towards increased noise and lower resolution.

### Weight of input sequence

To assess the influence of each sequence on our ICV and ventricular models by replacing each sequence with a highly noisy version (with a standard deviation of 0.5) in turn (**Suppl. Figure 2** and **Suppl. Figure 3**, respectively). The mean Dice coefficient for each noisy sequence is summarized in **Suppl. Table 1**. Lower Dice coefficients (on the validation dataset) were indicative of a greater weight or effect on the segmentation result. Running the ICV model with highly noisy inputs resulted in average Dice coefficients of 0.982 for the T1, 0.983 for the T2, and 0.984 for the FLAIR. While the Dice coefficient dropped across all sequences when segmenting the ICV, no noisy input sequence resulted in a substantial drop; hence these different sequences (contrasts) had similar weight on the segmentation, with T1 showing the largest effect. For the ventricular segmentation model, training on a very noisy T1 resulted in a significant drop in segmentation accuracy, with a Dice coefficient of 0.892, highlighting its importance for this segmentation task. The use of highly noisy T2 and FLAIR did not have a significant impact on Dice. Given the results of sequence impact on segmentation, we wanted to explore the segmentation results for models trained on different combinations of sequences for both regions of interest (ROIs) (**Suppl. Table 2**). While all trained model combinations produced a Dice coefficient overlap greater than 0.9, the models trained using three input sequences produced the highest Dice coefficients for both regions of interest, as expected.

**Figure 2.**
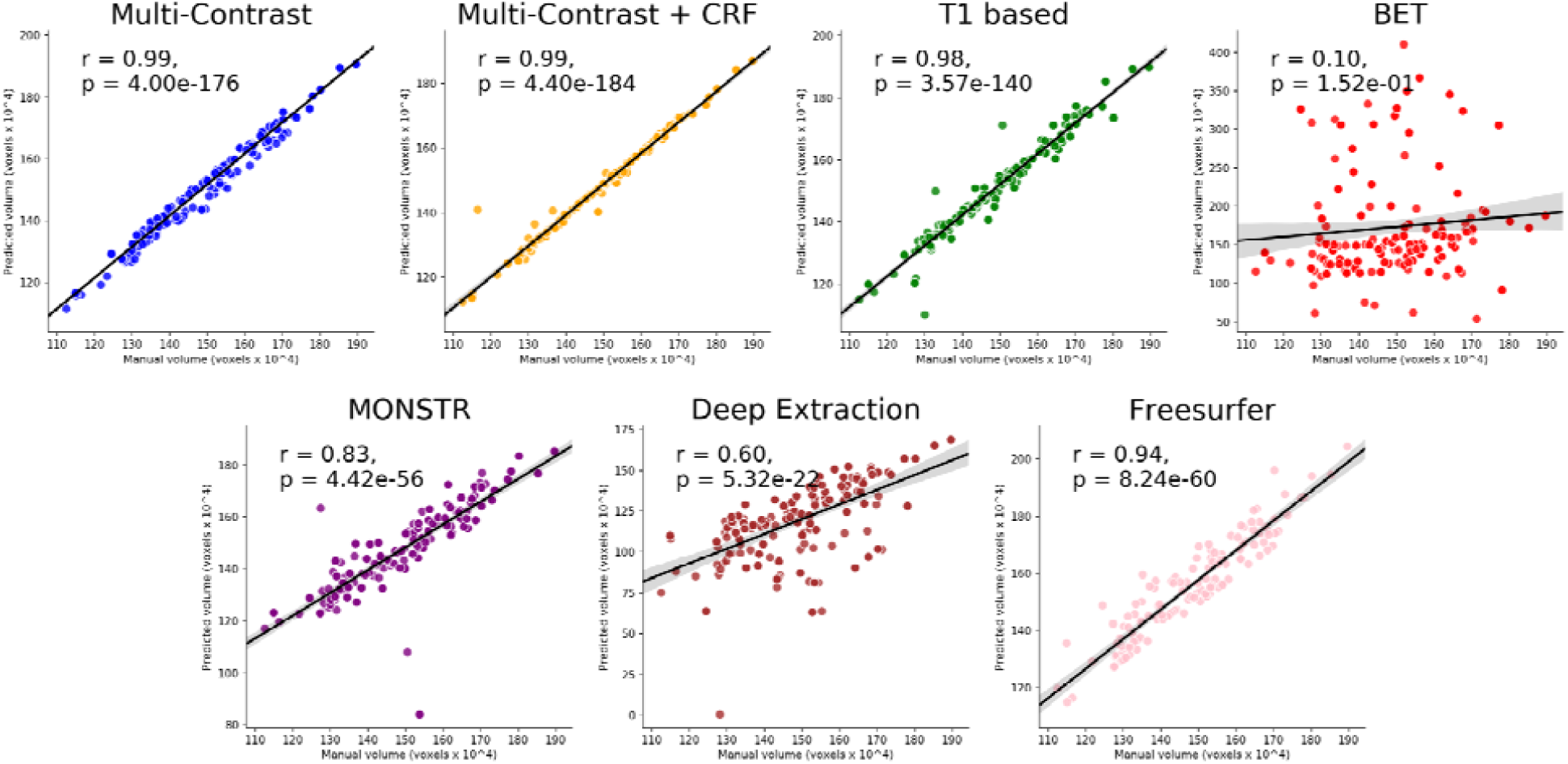
Pearson R. correlation coefficients and p-values for ICV segmentation between ground truth volumes and predicted volumes using our proposed multi-contrast model (iCVMapper, blue), multi-contrast plus CRF (yellow), our T1-based model (green), as well as four established techniques: MONSTR (purple), Deep Extraction (brown), FreeSurfer (pink) and BET (red).

### Evaluation of clinical datasets

#### Total Intracranial Volume Segmentation

Pearson correlations between the manual segmentation volumes and volumes predicted by the six tested methods are summarized in **Figure 2**. Our multi-contrast model had the highest volume correlations with manual ICV compared to other tested techniques (r=0.99, p<0.0001). The T1-based model and the multi-contrast-based MONSTR technique performed similarly based on volume correlations (r=0.84, p<0.0001), second best after our multi-contrast model, followed by FreeSurfer, MONSTR, Deep Extraction, and BET. BET did not produce significant correlations with manual ICV and had the lowest Dice and Jaccard score of all the methods.

Our multi-contrast model also performed the best among tested multi-contrast techniques (p < 0.001) on both the Dice and Jaccard coefficients (0.985 ± 0.01 and 0.971 ± 0.01 respectively, **Table 2** and **Figure 3**). Both the multi-contrast and T1-based models had a low Hausdorff distance (1.711 ± 0.481 mm for the multi-contrast model, 2.997 ± 2.726 mm for the T1-based model). MONSTR also performed well on overlap measures, with a Dice of 0.957 ± 0.080 and a Hausdorff distance of 5.733 ± 9.319 mm. BET’s box plot distributions across all metrics, as seen in **Figure 3**, reveal a large degree of segmentation variance, potentially a result of the inclusion of patients with lesions or high WMH burden in our test cohorts. When assessing the evaluation metrics across multi-contrast methods for the subset of patients with high WMH burden (a Fazekas score bigger than 2, from MITNEC study), we found that while all other SOTA methods had a relatively high Dice coefficient (> 0.80, our model’s Dice: 0.985), they had 3 to 12 times higher Hausdorff values than our multi-contrast model. For example, our model had an average Hausdorff value of 2.499 ± 4.632 mm while Deep Extraction’s Hausdorff value was 32.324 ± 13.014 mm. Similarly, other multi-contrast methods had 3 to 25 times higher AVD values on ICV segmentation of patients with WMH compared to our model (for example, our model’s AVD: 1.288 ± 2.95 and BET: 33.048 ± 40.107). While the T1-based model significantly outperformed FreeSurfer and Deep Extraction on the entire test set across all metrics (p < 0.001), outliers seen in the box plot of Hausdorff distances indicates the need for further fine tuning of model weights. Both the multi-contrast and T1-based models had higher Fischer z values (2.357 and 2.120, respectively) than each of their counterparts, indicating a closer correlation between their voxel counts and those of the ground truth.

To test the model’s performance with the CRF, we ran the CRF with five different parameter configurations on the test dataset and compared the resulting segmentations to the ground truth (**Suppl. Table 3**). Adjusting the parameters of the CRF to use smaller smoothness and appearance kernels resulted in a higher Dice coefficient than the multi-contrast model (0.99 ± 0.01); suggesting that using smaller neighbourhoods of voxels with similar intensity could aid in this segmentation task, probably due to MR image inhomogeneity. While the application of the CRF improved the average overlap and Dice coefficient, it resulted in a higher average Hausdorff distance and more outliers than the original model (**Figure 3**). All resulting segmentations using other CRF parameters had average Dice coefficients that were higher than all SOTA methods besides FreeSurfer.

**Figure 3.**
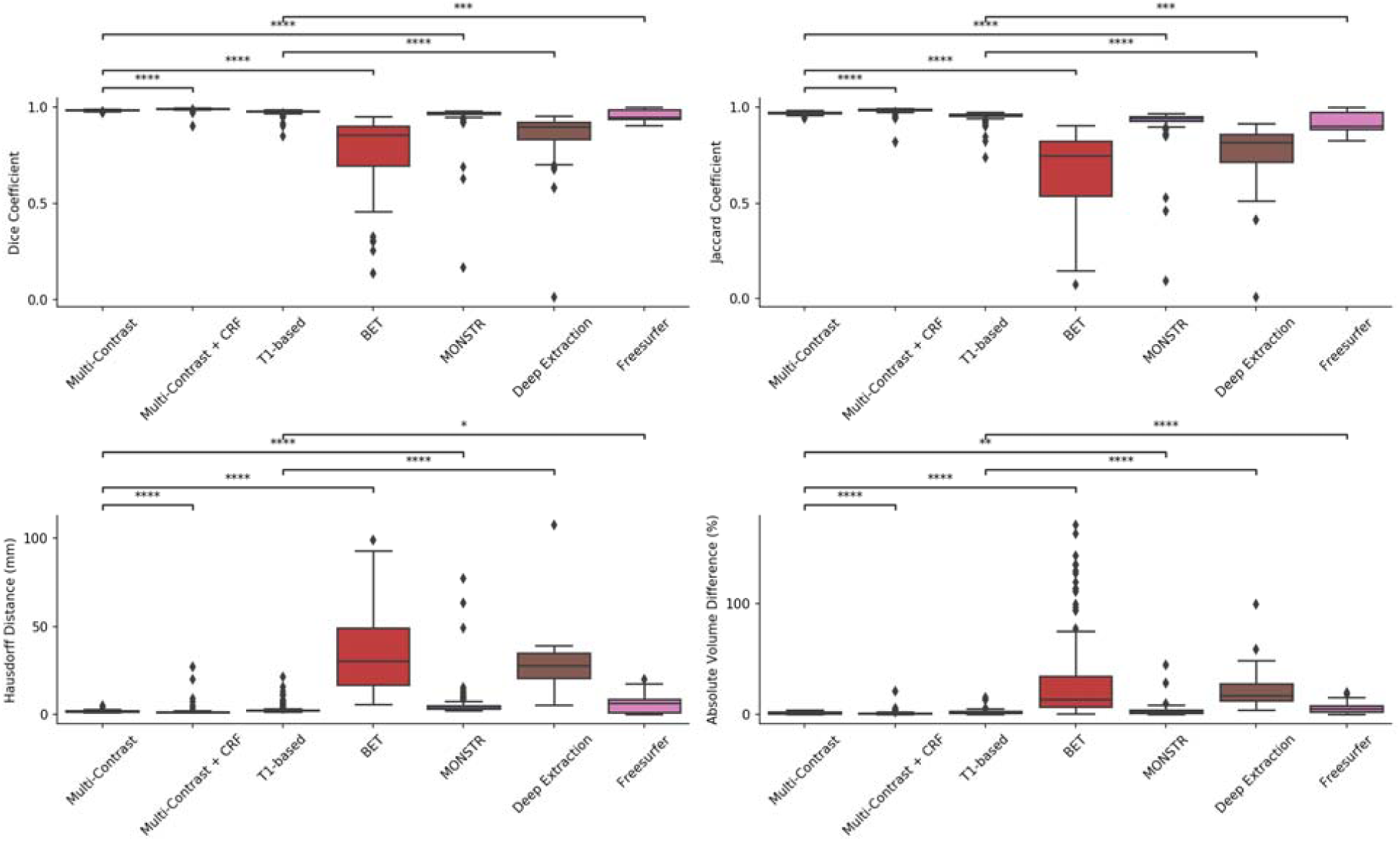
Dice and Jaccard coefficients, Hausdorff distance, and absolute volume difference between ground truth labels and predicted segmentations of the ICV. not significant: ns, p < 0.05: *; p < 0.01: **; p < 0.001: ***; p < 0.0001: ****.

The cases with the highest and lowest Dice coefficients between the manual brain segmentations and the multi-contrast model’s predictions are displayed in **Figure 4** (A and B, respectively). Mis-segmentations occurred more often in the dorsal slices, where distinguishing between tissue and dura may be difficult. The yellow arrows in **Figure 4A** highlight voxels that were falsely labeled as tissue. Visual inspection of the worst case demonstrates some of the challenges of the segmentation task, such as under-segmentation of the posterior regions, delineating the dura mater in the central fissure, and over-segmentation into the sinuses in subjects with atrophy. To appreciate the heterogeneity in our sample as a result of disease, we demonstrate the segmentation results from four individual test cases with brain lesions (**Suppl. Figure 3**). For example, the model was able to successfully segment the ICV up to the dura, even in the presence of strokes, and a large degree of atrophy (**Suppl. Figure 3A**). In some cases with brain lesions, mis-segmented voxels occurred around challenging anatomical regions at the superior and inferior edges of the cerebral cortex (intersection between brain parenchyma with subarachnoid space and dura), including the posterior longitudinal fissure and the pituitary gland (**Suppl. Figure 3B**).

**Figure 4.**
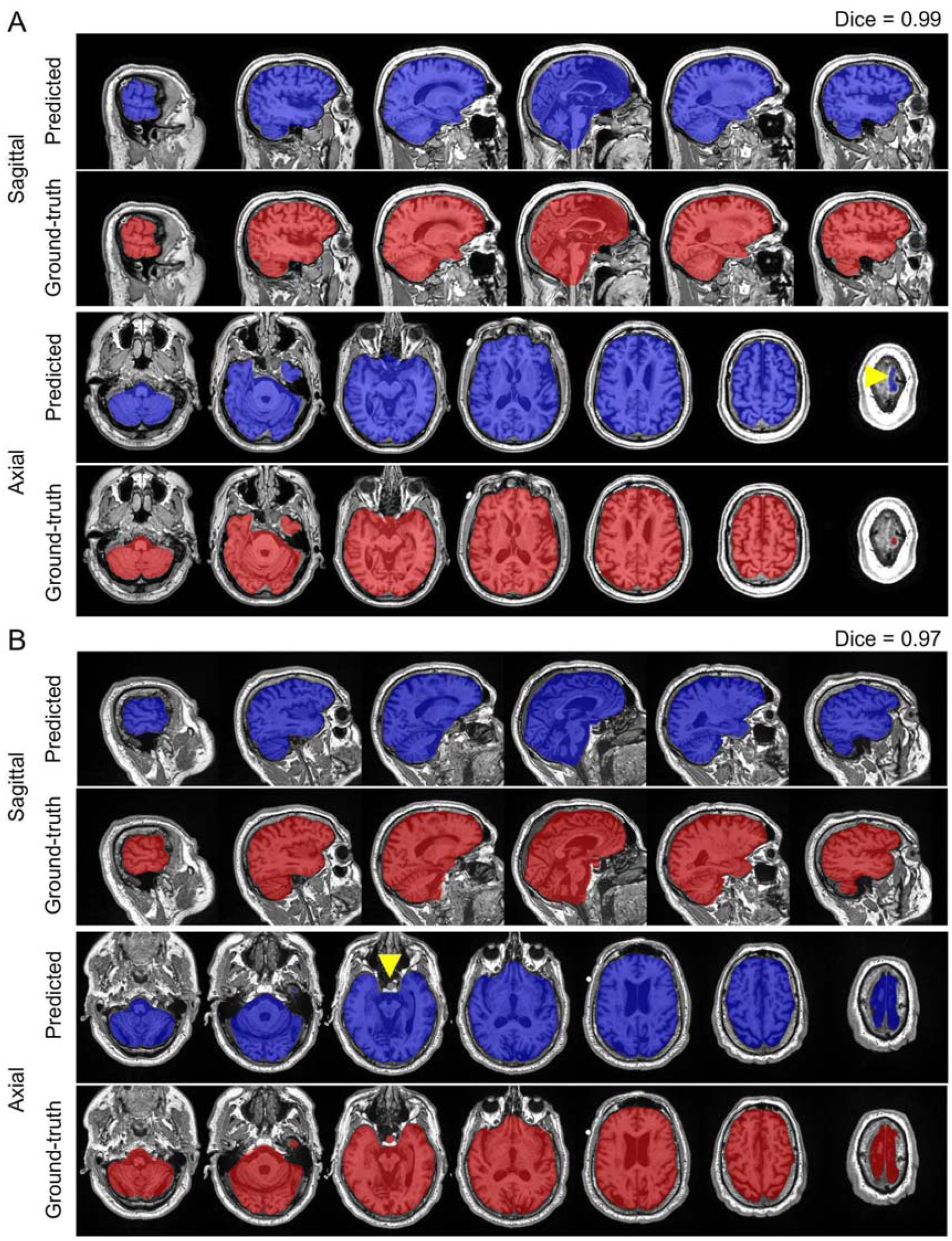
Skull stripping segmentation cases with the highest (A) and lowest (B) dice coefficients from the test set in the coronal and axial views. Blue labels represent predicted segmentations and red represent ground truth delineations. Yellow arrowheads highlight areas of miss-segmentations.

Each segmentation model was also tested on a cohort of patients diagnosed with VCI from the SDS, consisting of only T1-weighted scans. We separated our SDS analysis from the remaining test set to validate the robustness of our tools on a patient population with vascular pathology and lesions, and since only the T1-based model was used. The T1-based iCVMapper had significantly higher average Dice and Jaccard coefficients (p < 0.001), and lower AVD values (p < 0.001), compared to SOTA methods on SDS, including those using multi-contrast inputs (**Figure 5**). iCVMapper had a higher average Hausdorff distance than FreeSurfer (15.463 ± 10.853 and 8.024 ± 1.258, respectively) indicating the existence of some outliers with high surface distances from the ground truth. BET was not included in the ICV analysis as its outputs were manually corrected to generate the ground truth masks.

**Figure 5.**
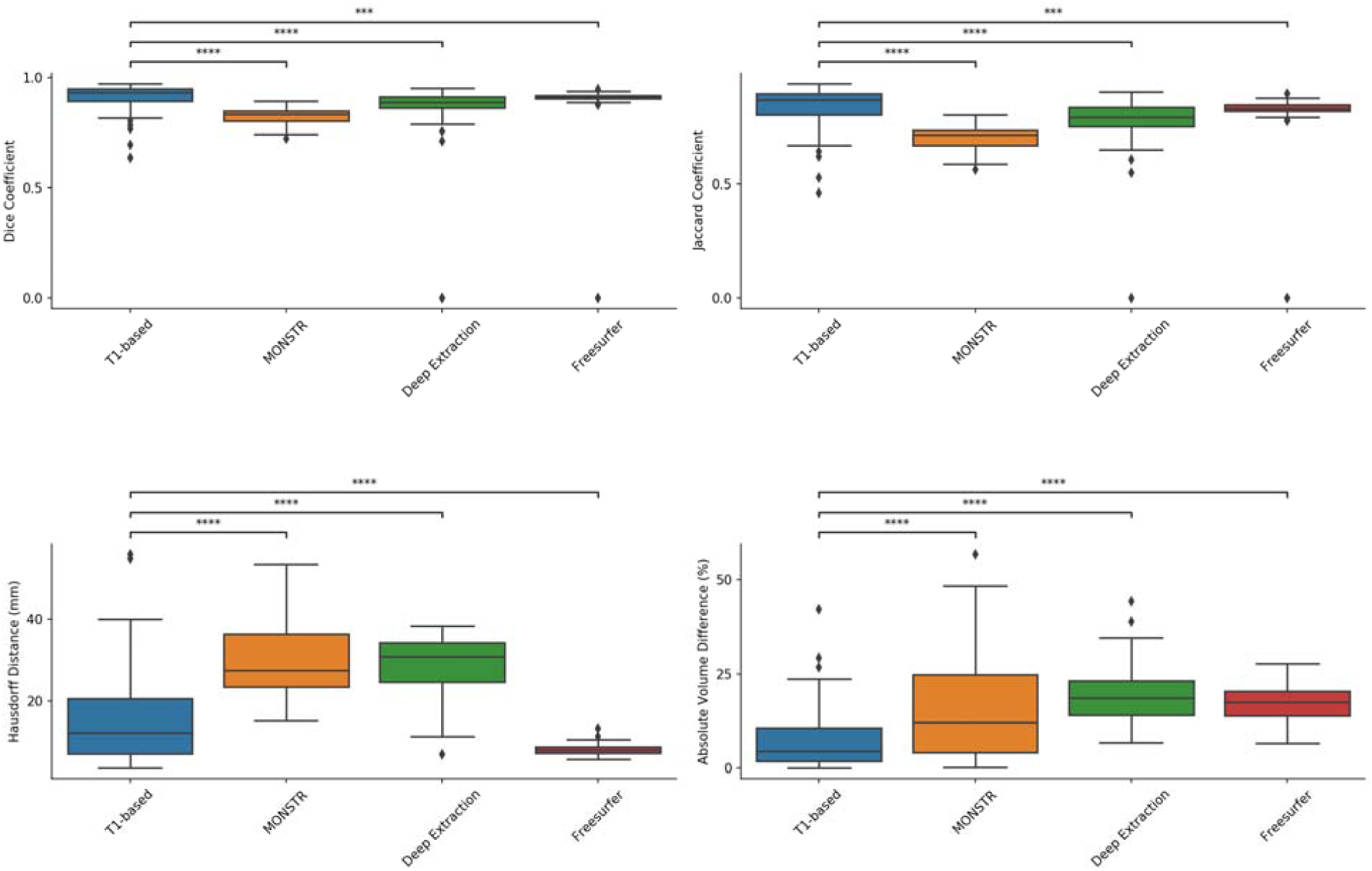
Boxplots of the distributions of Dice and Jaccard coefficients, Hausdorff distances, and absolute volume differences when segmenting the ICV of the VCI cohort of the Sunnybrook Dementia Study. not significant: ns, p < 0.05: *; p < 0.01: **; p < 0.001: ***; p < 0.0001: ****.

The computational time required for each method on a single core is also summarized in **Table 2**. Both the multi-contrast model and T1-based models had faster computational times than other methods, segmenting up to 300x faster than the slowest method. While Deep Extraction (the other tested CNN implementation) did not perform as well as other atlas-based algorithms such as FreeSurfer and MONSTR, it segmented the brain in under a minute.

**Table 2.** Dice, Jaccard coefficients, Hausdorff distance, absolute volume difference, Fisher’s z-transform, and computational time for ICV segmentation methods

|  | **iCVMapper**<br>(Multi-contrast) | **iCVMapper + CRF**<br>(Multi-contrast) | **iCVMapper**<br>(T1w only) | **MONSTR**<br>(Multi-contrast) | **BET**<br>(Multi-contrast) | **Deep Extraction**<br>(T1w only) | **FreeSurfer**<br>*<br>(T1w only) |
| --- | --- | --- | --- | --- | --- | --- | --- |
| **Dice coefficient** | 0.985 ± 0.003 | **0.993 ± 0.008** | 0.976 ± 0.016 | 0.957 ± 0.080 | 0.790 ± 0.166 | 0.863 ± 0.107 | 0.960 ± 0.027 |
| **Jaccard coefficient** | 0.970 ± 0.006 | **0.986 ± 0.014** | 0.954 ± 0.028 | 0.925 ± 0.094 | 0.679 ± 0.196 | 0.771 ± 0.129 | 0.924 ± 0.050 |
| **Hausdorff Distance (mm)** | **1.711 ± 0.481** | 1.836 ± 2.908 | 2.997 ± 2.726 | 5.733 ± 9.319 | 36.217 ± 24.743 | 27.155 ± 11.507 | 5.660 ± 4.558 |
| **Absolute Volume Difference (%)** | 1.591 ± 0.962 | **1.088 ± 1.894** | 2.224 ± 2.204 | 3.295 ± 5.290 | 30.569 ± 39.551 | 21.418 ± 13.677 | 5.210 ± 4.294 |
| **Fisher’s Z** | 2.357 | **2.513** | 2.120 | 1.241 | 0.091 | 0.680 | 1.721 |
| **Approx. computational time** | 11 s | 4 min | **9 s** | 45 min<br>(3 contrasts, 6 atlases) | 3 min | 43 s | 20.7 min |

\*FreeSurfer failed to process three scans.

#### Ventricular Segmentation

All tested methods for ventricular segmentation had significant agreement between their respective predicted segmentation volume and manual segmentation volumes, as indicated by the high correlation coefficients (**Figure 6**). Average Dice and Jaccard coefficients, Hausdorff distances, and absolute volume differences for VentMapper and FreeSurfer ventricular segmentations are summarized in **Table 3** and **Figure 7**. Both the multi-contrast model and the T1-based network had higher Dice coefficients (0.945 ± 0.022 and 0.945 ± 0.021, respectively) than FreeSurfer (0.90 ± 0.04), and lower Hausdorff scores. This was apparent in the ability of both networks to segment areas of high difficulty, such as disjoint temporal horns. While the T1-based model had a lower average Dice and Jaccard coefficient relative to the multi-contrast model paired with the CRF, the Hausdorff distance was marginally lower. Both the multi-contrast and T1-based method algorithms were two orders of magnitude faster than that of FreeSurfer. This is due to the fact that FreeSurfer performs surface and image-based registrations and segmentations of many brain structures other than the ventricles and thus is more computationally demanding. FreeSurfer’s failed cases (incomplete segmentation pipeline) were excluded from the analysis.

**Table 3.**
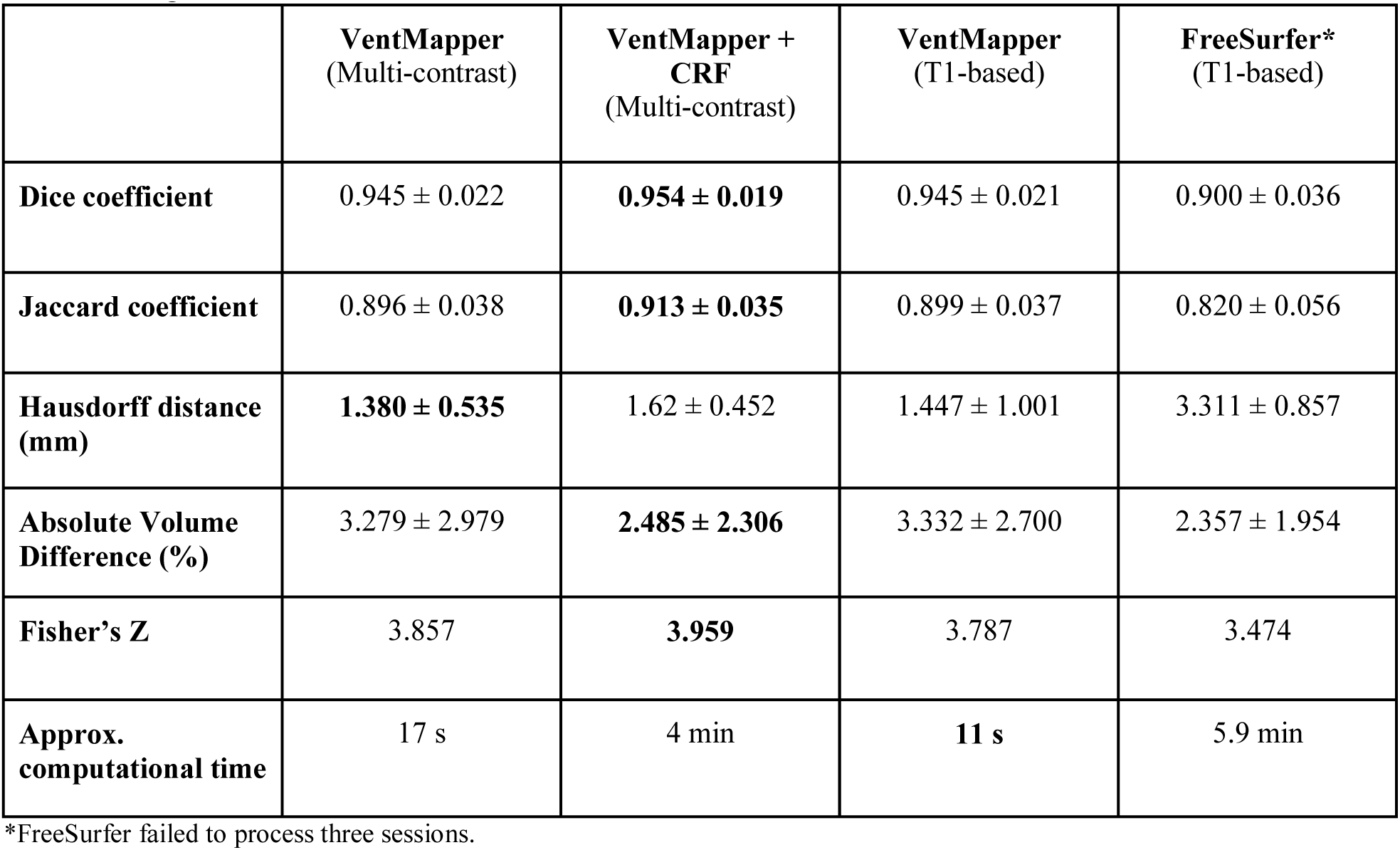
Dice, Jaccard coefficients, Hausdorff distance, absolute volume difference, and computational time for ventricular segmentation methods

**Figure 6.**
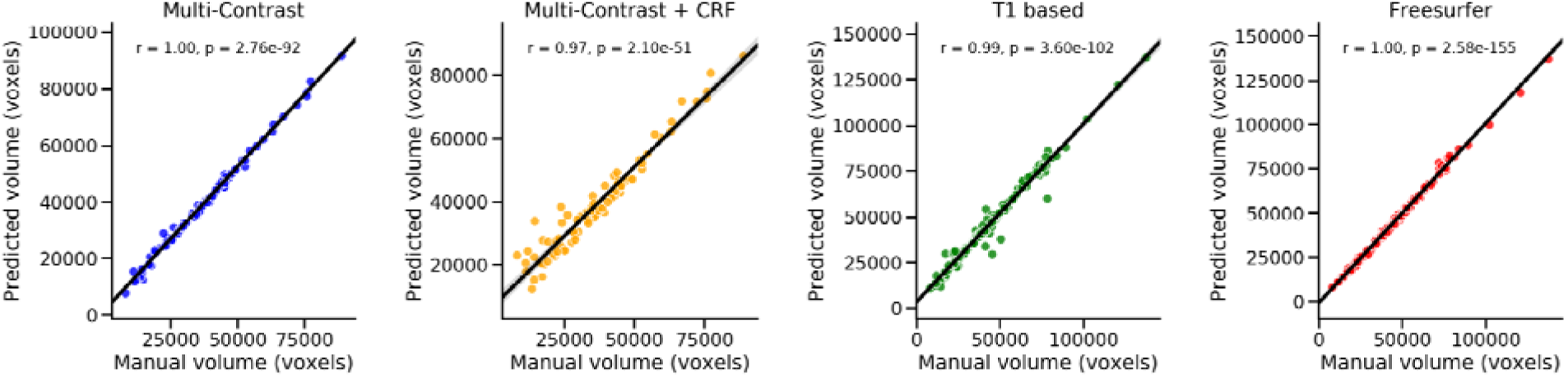
Pearson R. correlation coefficients and p-values between the manually segmented volumes and volumes generated through model predictions.

**Figure 7.**
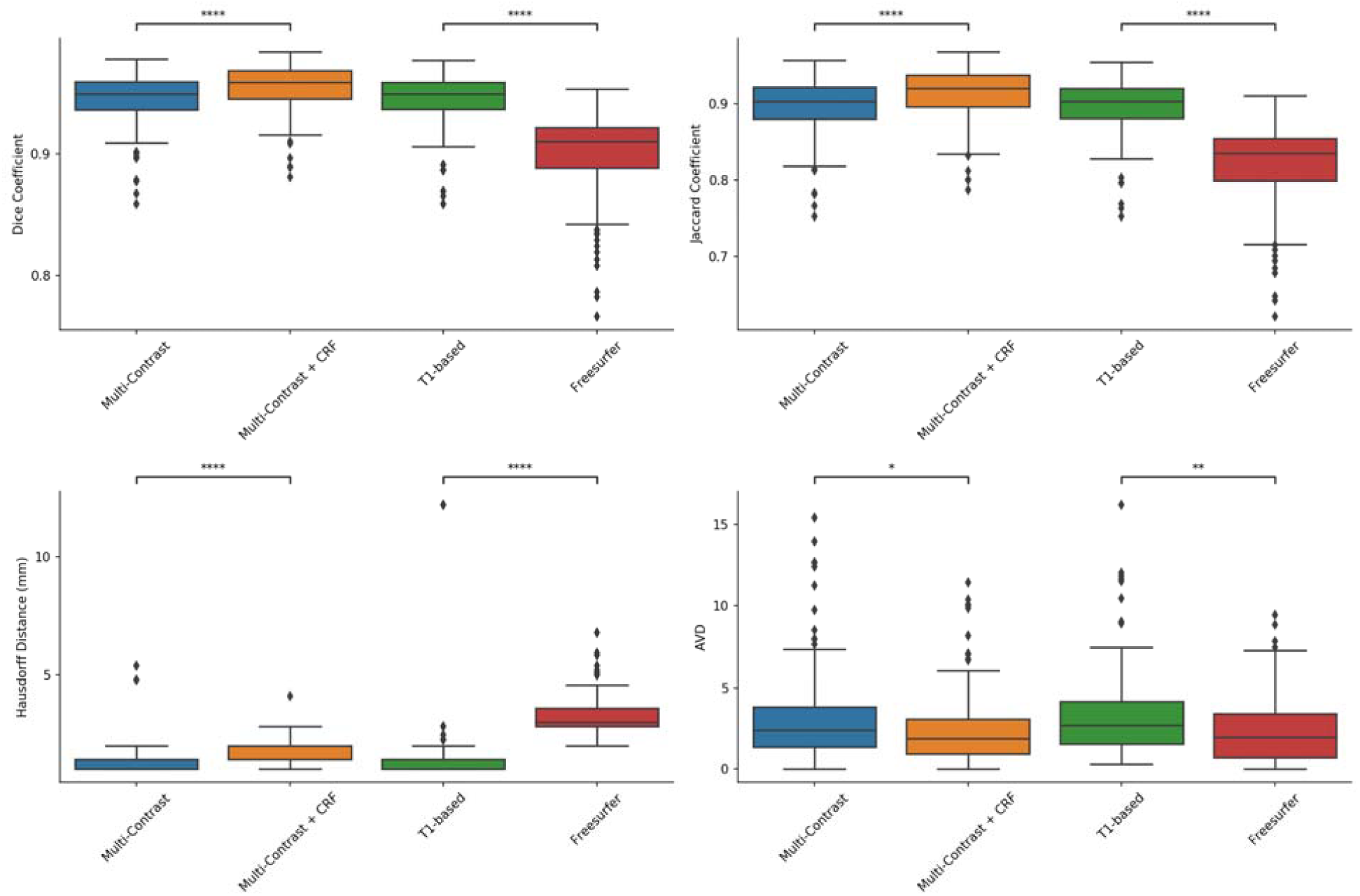
Dice and Jaccard coefficients, Hausdorff distance, and absolute volume difference (AVD) between ground truth labels and predicted segmentations of the ventricles. not significant: ns, p < 0.05: *; p < 0.01: **; p < 0.001: ***; p < 0.0001: ****.

VentMapper’s T1-based model’s segmentation results were compared to that of FreeSurfer on SDS’s VCI cohort. VentMapper had a better performance than FreeSurfer across all four metrics (**Suppl. Figure 5**), notably having Dice coefficient of 0.951 ± 0.021 (p < 0.0001, Mann-Whitney *U* two-tailed test) and a Hausdorff distance of 1.881 ± 2.278 mm (p < 0.0001, Mann-Whitney *U* two-tailed). While the Hausdorff distance for VentMapper was lower (p < 0.0001), outliers were found for both VentMapper and FreeSurfer’s segmentations. This observation, paired with the outliers found in the AVD box plot may indicate that there were sections of tissue that were misclassified as ventricular CSF, rather than false positive voxels.

To test the model’s performance with the CRF, the same parameter configurations that were used on the ICV segmentation dataset were applied to the ventricular segmentation results (**Suppl. Table 4**). Two of the five combinations that were tested improved the performance of the multi-contrast model. The model with the lowest smoothness and appearance kernel metrics resulted in the greatest improvement, producing an average Dice coefficient of 0.954 ± 0.019. This may indicate that conservative kernels are more optimal when applying post processing to high performing segmentation models. On average, the Hausdorff distance for the CRF was greater than that of the multi-contrast model, despite having a better performance across other metrics. It is possible that this is a result of poor smoothing as a result of the kernel parameters utilized. A more exhaustive exploration of the parameters of the appearance kernel may be needed to produce an improved ventricular segmentation.

The two cases with highest and lowest Dice coefficients using our multi-contrast network are shown in **Figure 8**. In both cases, the model was able to segment the ventricles within anatomical regions with high complexity, such as the occipital horns. To highlight our ventricular network robustness, we demonstrate our multi-contrast model results on four challenging test cases with strokes, ventricular enlargement, brain atrophy, and low SNR in **Figure 11** and **Suppl. Fig. 4**. The multi-contrast model was able to efficiently distinguish between the ventricles and diffuse periventricular WMH, as indicated by the yellow arrows. Cases with high Hausdorff distances tended to miss extensions of the ventricles that thinned as they moved into the temporal lobes. This may be due to downsampling of the input images during training.

**Figure 8.**
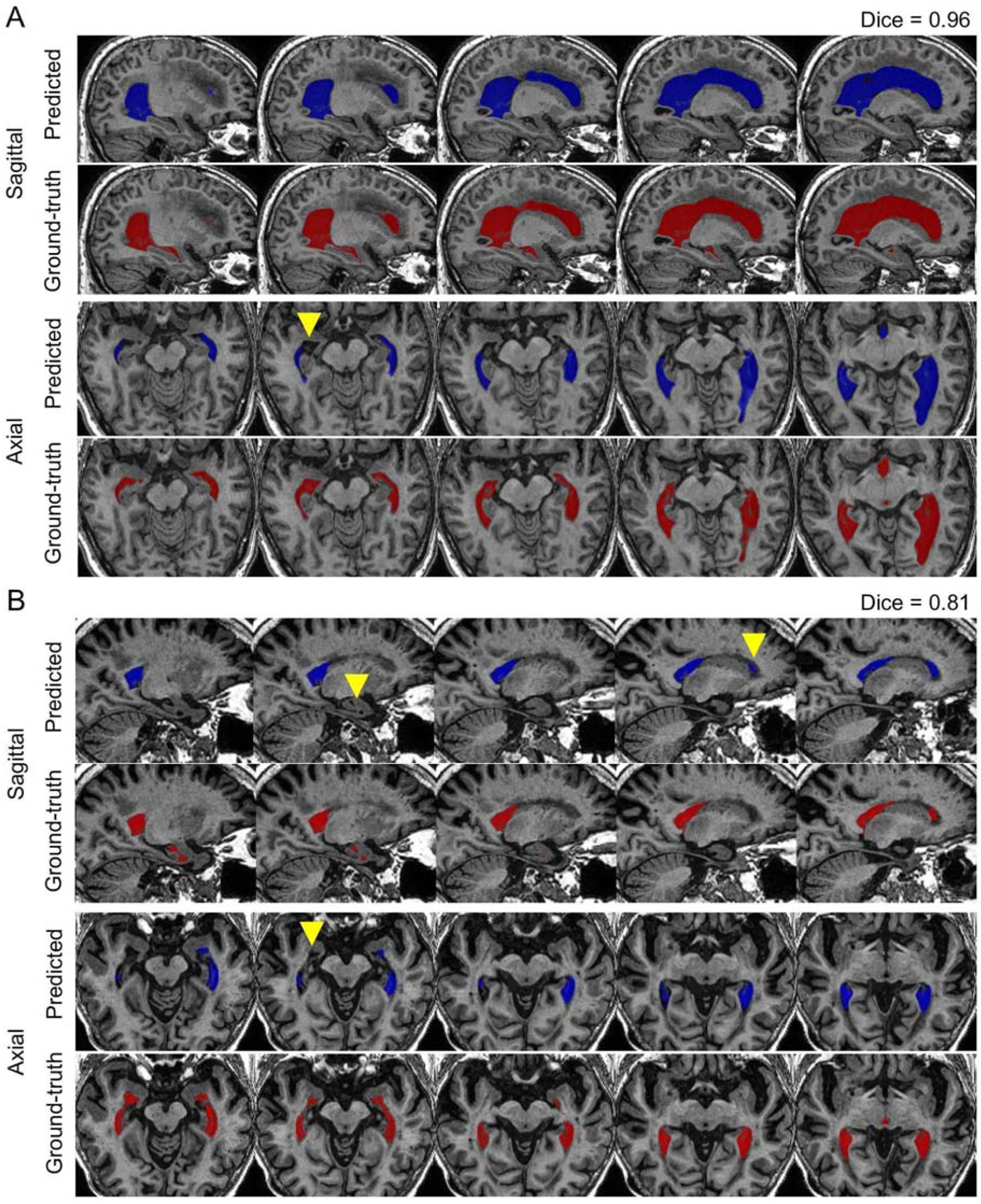
Ventricular segmentation cases with the highest (A) and lowest (B) Dice coefficients from the test set in the sagittal and axial views. Blue labels represent predicted segmentations and red labels represent manual delineations. Yellow arrowheads highlight areas of missed segmentations.

**Figure 9.**
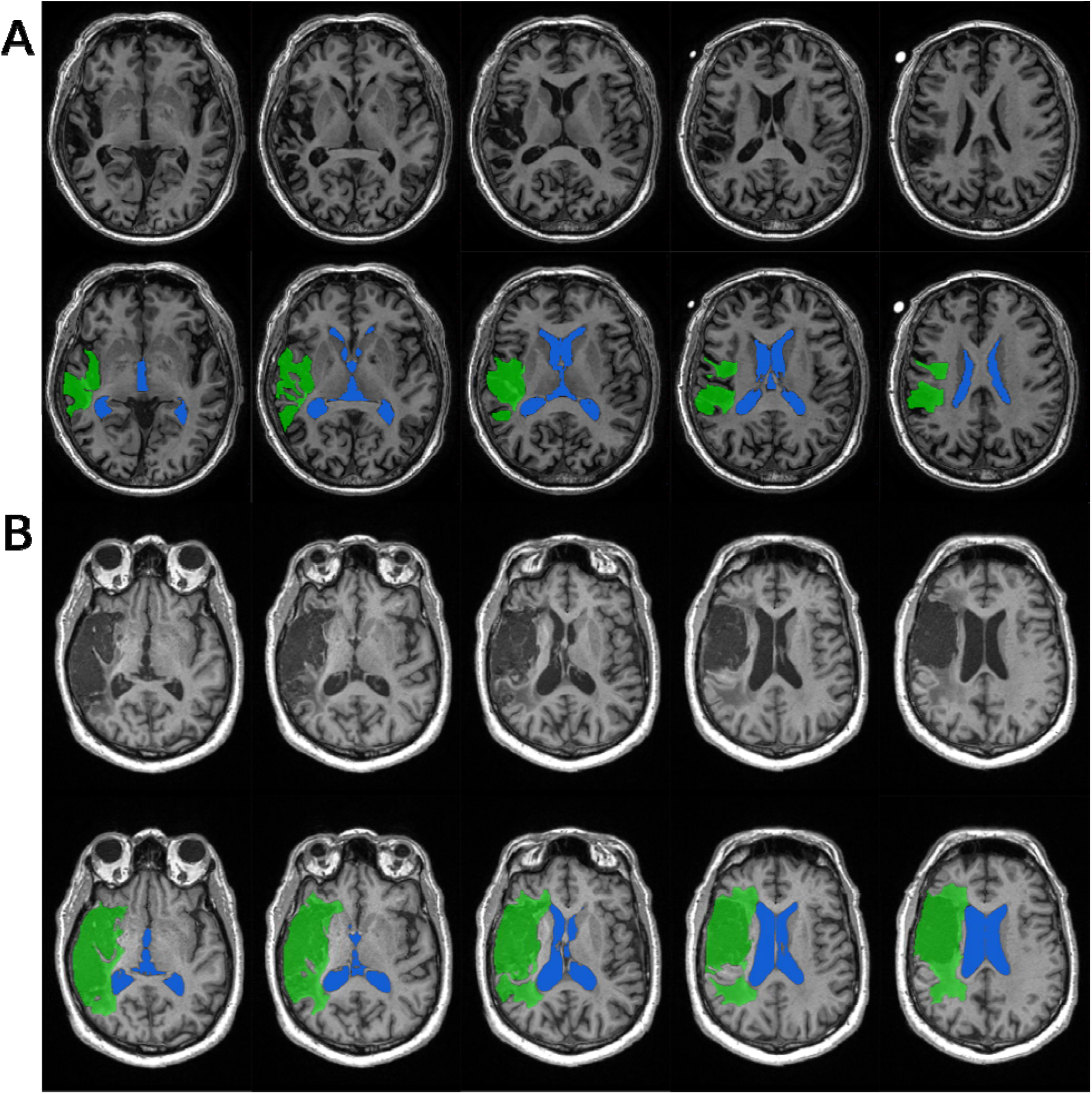
Ventricular segmentation (indicated by blue label) in two cases (A and B) with large ischemic strokes (green labels) adjacent to the ventricles, demonstrating the accuracy of our model in these challenging cases

**Figure 10.**
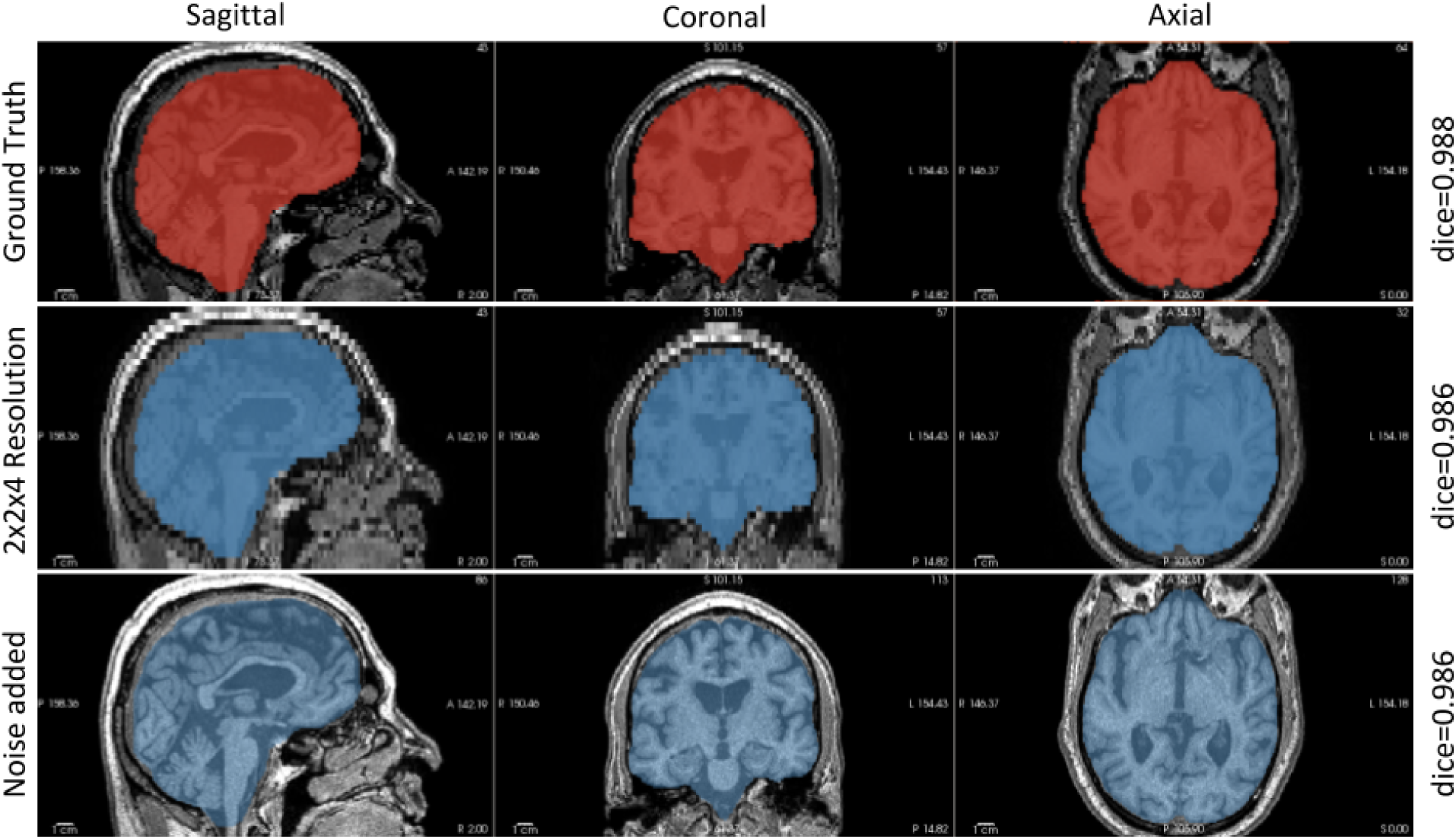
Brain segmentation in two cases of adversarial attacks (downsampling of resolution by a factor of 2×2×4, and the addition of gamma noise with a sigma of 0.1) applied to the same image.

**Figure 11.**
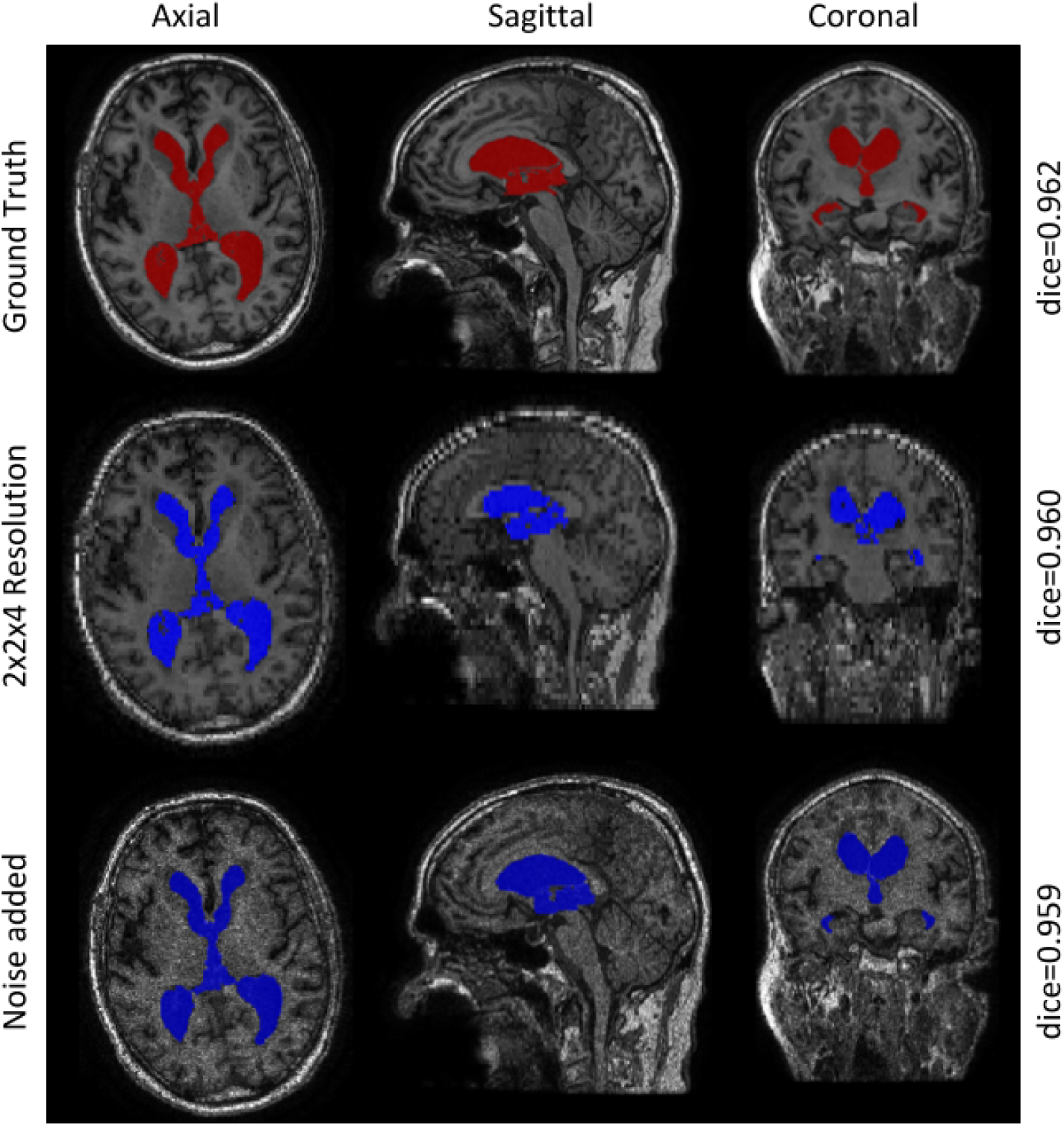
Ventricular segmentation in two cases of adversarial attacks (downsampling of resolution by a factor of 2×2×4, and the addition of multiplicative noise with a sigma of 0.1) applied to the same image.

#### Clinical adversarial cases

The effects of low resolution and signal-to-noise ratio on our models’ performance were simulated by testing both models on downsampled and noisy images. ICV and ventricular segmentation results for these simulated images are shown in **Figure 10** and **Figure 11**, respectively. For both the ICV and ventricular segmentations there was no substantial decline in the multi-contrast model’s performance, regardless of the downsampling factor (**Suppl. Table 5**). For both tested voxel sizes, iCVMapper produced an overall marginal drop of less than 0.2% when employing the three contrasts, while the T1-based segmentation model resulted in an average drop of 4.7%. VentMapper had a similar performance across both downsampling experiments, with average drops in the Dice coefficient ranging between 1.7 and 2.4%. The small percent reductions in Dice coefficient demonstrate both models’ ability to produce accurate segmentations when encountering acquisitions with lower resolutions.

Increases in simulated noise correlated with an increase in overall Dice coefficient drop for both the ICV and ventricular segmentation models. The T1-based model showed a greater overall decrease in Dice coefficient (3.0%) relative to the multi-contrast model (0.1%). Over-segmentation occurred in superior and posterior regions. Similarly, for the ventricular segmentation with the highest tested noise sigma, the multi-contrast model produced a slightly higher Dice drop (3.8%) relative to our T1-based model (3.1%). The multi-contrast model had a more substantial drop in Dice for ventricular segmentation with noisy inputs as compared to brain segmentation, which was similar in magnitude to the T1-based mode. This highlights the higher complexity of the task and the segmentation model’s potential difficulty in extracting unique features from different contrasts when relying on input data with low SNR.

All SOTA methods were compared to our models on both downsampled by a factor of 2, 2 and 4 across the respective x, y and z planes, as well as the noisy data by a sigma of 0.5 (**Suppl Tables 6, 7, 8** and **9**). The majority of cases failed for both adversarial attacks when employing Deep Extraction, hence it was not included in this comparison. FreeSurfer failed to generate 7 ICV and ventricular segmentation masks for downsampled adversarial cases, and 42 (out of 132) ICV and ventricular segmentation for noise induced cases. MONSTR failed to segment 5 of the noisy cases. Failed cases were not included in the analysis or generated figures. The adversarial attacks did not lead to substantial changes in the computed volumes for both ICV and Ventricular segmentation using our models; however, we found that the artificial adversarial cases had significant effects on the other SOTA methods (except for FreeSurfer on downsampled cases) as apparent by the substantial increases in AVD values (**Suppl Tables 6, 7, 8** and **9**).

For ICV segmentation on adversarial cases with lower resolution, all other SOTA methods besides FreeSurfer had significant decreases in performance as a result of downsampling the images. The multi-contrast version of iCVMapper had a better performance on the Dice and Jaccard coefficients, and Hausdorff distance when compared to BET and MONSTR (**Figure 12**). Notably, MONSTR had a lower AVD than our multi-contrast model. The T1-based iCVMapper had a Dice coefficient of 0.866 ± 0.096 and a Hausdorff distance of 2.896 ± 3.522 mm, which was better than Deep Extraction but worse than FreeSurfer when excluding failed cases (Dice coefficient: 0.928 ± 0.019, Hausdorff distance: 4.685 ± 1.300 mm). For ventricular segmentation of downsampled cases, T1-based VentMapper outperformed FreeSurfer on Dice and Jaccard coefficients and Hausdorff distance, but had a higher AVD (7.659 ± 6.150 vs. 2.954 ± 2.410). This would indicate that while the absolute number of voxels classified as ventricles was closer to that of the ground truth in FreeSurfer, many of these voxels were misclassified false positive with a significantly worse overlap and average surface distance than VentMapper.

**Figure 12.**
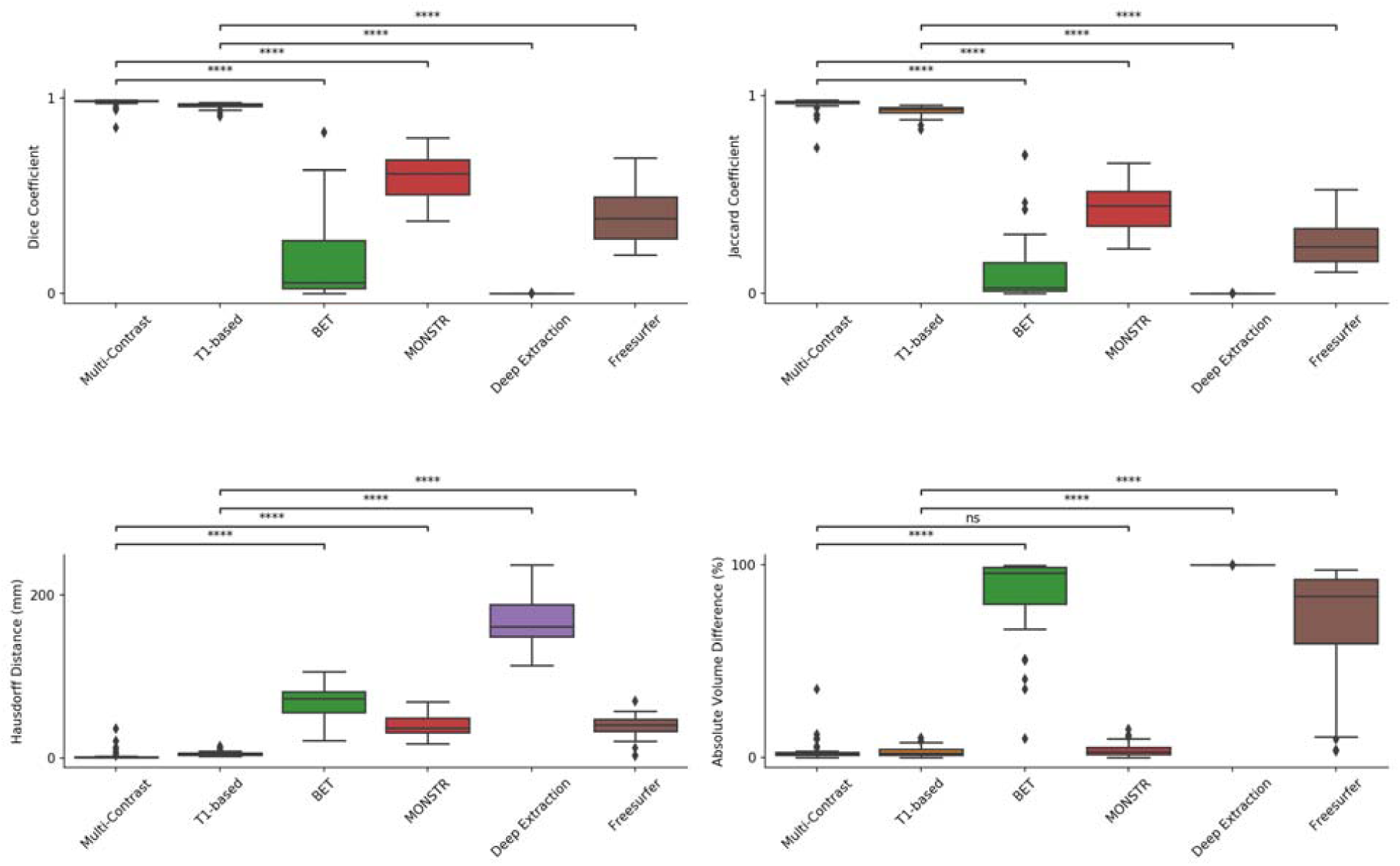
Dice and Jaccard coefficients, Hausdorff distance, and absolute volume difference of ICV segmentation on adversarial cases with increased noise. not significant: ns, p < 0.05: *; p < 0.01: **; p < 0.001: ***; p < 0.0001: ****.

The iCVMapper and VentMapper had better performances across all metrics for the noise induced cases in comparison to the other segmentation methods. The results are demonstrated in **Supplementary Figures 5 and 6**. As indicated by the aforementioned low drop in Dice coefficient, ICVMapper’s multi-contrast and T1-based models had significantly lower AVDs relative to other SOTA methods (multi-contrast: 2.235 ± 3.47, T1 based: 3.626 ± 3.551). The T1-based model had a higher Hausdorff distance than the multi-contrast model (6.044 ± 4.307 mm vs 2.907 ± 4.321 mm). Comparing the VentMapper’s T1-based model and FreeSurfer, we found that VentMapper’s Dice coefficient and Hausdorff distance (0.909 ± 0.036 and 1.785 ± 0.759 mm) indicated a significantly improved overlap and segmentation fidelity relative to FreeSurfer (Dice coefficient: 0.154 ± 0.247, Hausdorff distance: 40.287 ± 15.821 mm). However, FreeSurfer’s ventricular segmentation could be attributed to its poor ICV segmentation (or initialization) which had a Dice coefficient of 0.386 ± 0.149. The results indicate that iCVMapper and VentMapper are robust to drops in signal-to-noise ratio of input data for these segmentation tasks, but more fine tuning may be required to improve the model’s performance segmenting ventricles on input data with lower resolutions.

## Discussion

We developed segmentation models for important structural biomarkers (total intracranial and ventricular volumes), that are robust to challenges that commonly result in segmentation failures. Our models were trained using heterogeneous multi-site, multi-scanner datasets with increased gray/white atrophy, larger ventricles, infarcts, lacunes, and white matter hyperintensities, reflective of an aging population. First, we assessed the sequence effects on segmentation accuracy to study CNN sensitivity to image weighting in our application. Next, we compared our algorithms to available SOTA methods demonstrating improvements in both accuracy and speed on test data characterized by vascular pathology and increased white matter disease burden from multiple studies (two of which were not used for training). We then tested various iterations of a CRF as a post-processing step to improve segmentation results. Finally, to simulate variability in image quality in clinical image acquisition we tested our models under the stressors of downsampling and lower SNR.

While numerous segmentation algorithms have been developed to segment the ICV and ventricles, the majority of available methods struggle in the face of vascular lesions and brain atrophy. Those that are robust to brain atrophy require parameter optimization or large computational run times and are thus complex to use for the naïve user and are not practical for large multi-site studies. Our models provide an open-source, accurate and efficient solution to segment these structures that require no parameter optimization and are easy to use; highlighting their applicability in large neuroimaging initiatives or cohorts. Our multi-contrast models achieved an average Dice overlap coefficient of 0.985 ± 0.003 and 0.945 ± 0.022 for brain and ventricular segmentations respectively in this difficult test dataset, the highest overlap coefficients among tested methods. The consistently high performance of our multi-contrast method highlights its robustness against different scan protocols and disease groups. They were orders of magnitude faster than the majority of other methods, segmenting the ICV and ventricles in seconds.

The Dice coefficient, Jaccard index, Hausdorff distance, absolute volume difference, and Pearson’s correlation coefficient were employed to quantitatively assess the performance of all segmentation methods. Although the absolute volume difference serves as a measurement of the similarity in voxel counts, it was included as a comparison metric to provide a fuller context to the segmentation results. While the Dice coefficient is commonly used to assess performance when segmenting images, it is less indicative of mismatch when segmenting larger ROIs, as is the case when segmenting the ICV. Thus, it should be reviewed in conjunction with surface distance metrics such as the Hausdorff distance. Assessing the Dice coefficient and Hausdorff scores for ICV segmentation, it is evident that our proposed models improve on the available SOTA methods. We found that using a CRF with the chosen parameters (**Suppl. Table 3**) improved the performance of both ICV and ventricular segmentations. Upon further inspection of the segmentation results, we observed that the CRF effectively removed false positives at the cost of eroding the segmentation result, which is due to the smoothing and appearance kernels (parameters) used for the CRF.

For ICV segmentation, MONSTR had the second highest Dice and second lowest Hausdorff but took on average 45 minutes longer to generate the segmentation. It uses a multi-atlas approach and computes similarity between the input subject and atlases along a thresholded band representing the outermost layer of the estimated brain mask (Roy et al. 2017). While the resultant segmentations are robust to various cases, the running time is contingent on the number of atlases and modalities utilized to obtain a segmented brain mask. In terms of speed, the algorithms based on CNNs (including ours) had processing times that were fast enough to be considered for use in a clinical setting. While the CNN-based Deep Extraction’s (Kleesiek et al. 2016) processing time was similar to our models, its lower accuracy on the test set may be due to the smaller number of subjects used for training. Since FreeSurfer’s segmentation and parcellation occurs in a sequential pipeline, their runtimes were extracted from the log files generated for each session for only that step. Besides FreeSurfer, all software that was compared to ICVMapper and VentMapper exclusively segmented the corresponding ROI.

We chose to train our model on segmentations that were semi-automatically edited by experts rather than automated outputs produced by other algorithms as they are expected to be more accurate. We further relied on data augmentation to provide more training datasets as commonly performed in training neural networks. We decided on a 3D architecture as it might better account for partial volume effects near the edges of the segmented brain and ventricles. Since 3D architectures are very computationally demanding, instance normalization (Ulyanov, Vedaldi, and Lempitsky 2016) was used to avoid having to use a large amount of data (in memory) in contrast to batch normalization.

Both a T1-based and a multiple contrast-based model were developed for each ROI to take advantage of different input sequences. Evaluating the differences in evaluation metrics between the multi-contrast and T1-based models, it is evident that accuracy improves with additional contrasts and hence additional important features are learned from the other sequences. This is of particular importance when accounting for brain lesions such as strokes and WMH, which may have different features under different MRI sequences.

To assess the robustness and generalizability of our models against inputs with lower quality than our training set, we simulated scans with lower resolution or SNR. These “adversarial cases” suggest that our multi-contrast models for both structures are robust to both attacks with varying degrees of intensity; whereas, the T1-based models are sensitive to downsampling as demonstrated by the decrease in Dice coefficient. This lends to the significance of utilizing multiple sequences or channels in segmentation tasks. While the T1-models were less sensitive to the addition of noise than downsampling, the ventricular network was affected by the introduction of noise with a high sigma. This finding cautions on the use of T1-based models in datasets with very low SNR.

Many of our conclusions were validated after the models’ performance segmenting the “adversarial cases” was compared to that of existing SOTA segmentation methods. SOTA methods that we compared against had widely varying results when segmenting the lower resolution and SNR data, with significant volume differences and substantial drops in quantitative metrics or segmentation fidelity, while our models remained largely robust to both simulated adversarial attacks. Notably, Deep Extraction failed on the majority of cases for both attacks, highlighting the sensitivity of some deep learning networks to out-of-distribution perturbations and the need for further validation of the generalizability of these networks. Further improvements of our models may also be necessary in light of some of the weaknesses found, most notably improving T1-based VentMapper’s performance on downsampled data. In addition to the attacks we simulated there are other challenges that could be tested in future work such as simulation of significant subject motion.

Though our model performed well compared to other segmentation algorithms, there are still some future improvements that can be made. While our multi-contrast model demonstrated robustness against adversarial cases, their use may be limited by a lack of standardization across imaging protocols, as different studies may not have the contrasts required as inputs. The models also have difficulty segmenting inputs that are not in a standard orientation. A potential solution would involve augmenting the data such that the inputs were flipped at a 90° angle across all three axes. Although we utilized large multi-site datasets encompassing several types of lesions and atrophies, our training data was still limited as it did not include other disorders such as normal pressure hydrocephalus or presence of different lesions such as brain tumors and brain contusions.

## Conclusions

We present robust and efficient segmentation methods for the ICV and ventricles, commonly used biomarkers of brain atrophy in normal aging and neurodegeneration. We trained CNN models with expert manually edited segmentations from large multi-site studies including participants with vascular lesions and atrophy, which represent challenging populations for segmentation techniques. Both our segmentation models achieved higher accuracy compared to state-of-the-art algorithms and were orders of magnitude faster than the majority of available methods. Additionally, our algorithms were robust to simulated images with low resolution and SNR. The pipeline and trained models are available at: https://icvmapp3r.readthedocs.io and https://ventmapp3r.readthedocs.io.

## Acknowledgements

We are grateful for the support of the Medical Imaging Trial Network of Canada (MITNEC) Grant #NCT02330510, and the following site Principal investigators: Christian Bocti, Michael Borrie, Howard Chertkow, Richard Frayne, Robin Hsiung, Robert Laforce, Jr., Michael D. Noseworthy, Frank S. Prato, Demetrios J. Sahlas, Eric E. Smith, Vesna Sossi, Alex Thiel, Jean-Paul Soucy, and Jean-Claude Tardif. We are also grateful for the support of the Canadian Atherosclerosis Imaging Network (CAIN) (http://www.canadianimagingnetwork.org/), and the following investigators: Therese Heinonen, Rob Beanlands, David Spence, Philippe L’Allier, Brian Rutt, Aaron Fenster, Matthias Friedrich, Ben Chow, and Richard Frayne. This research was conducted with the support of the Ontario Brain Institute, an independent non-profit corporation, funded partially by the Ontario government. The opinions, results and conclusions are those of the authors and no endorsement by the Ontario Brain Institute is intended or should be inferred.

## Declarations

### Data Accessibility

The developed algorithm and trained models (network weights) are publicly available at: https://icvmapp3r.readthedocs.io and https://ventmapp3r.readthedocs.io under the GNU General Public License v3.0. An example dataset is included for testing purposes. We have developed an easy□to□use pipeline with a GUI and thorough documentation for making it accessible to users without programming knowledge.

## SUPPLEMENTARY FIGURES

**Suppl. Figure 1**. Effect of input sequence (contrast) on ICV extraction accuracy. Blue represent overlap, Red voxels present in ground truth missing in prediction, Green voxels present in prediction and not in ground truth.

**Suppl. Figure 2**. Effect of input sequence (contrast) on ventricular segmentation accuracy. Blue represent overlap, Red voxels present in ground truth missing in prediction, Green voxels present in prediction and not in ground truth.

**Suppl. Figure 3**. Brain segmentation in four test cases from different studies highlighting the quality of segmentations. Yellow arrowheads show strokes.

(A) Predicted segmentations (blue label) overlaid on manual delineations (red outline).

(B) Miss-segmented voxels highlight the differences between the segmentations. Red voxels are manually labeled voxels not predicted by the model, while light blue voxels are predicted voxels that were not present in the manual labels.

**Suppl. Figure 4**. Ventricular segmentation in four test cases from different studies highlighting the quality of segmentations. Yellow arrowheads show severe white matter hyperintensities (WMH) or stroke.

(A) Shows under segmentation of our model (blue label) by overlay on manual delineations (red).

(B) Shows the overlap in our model compared to the manual segmentation by voxels incorrectly labelled by our model that were not in the manual segmentation (light blue) and voxel missed by our model that were present in the manual segmentation (red).

**Suppl. Fig. 5**. Dice and Jaccard coefficients, Hausdorff distance, and absolute volume difference of ventricular segmentation on the SDS vascular cognitive impairment cohort. not significant: ns, p < 0.05: *; p < 0.01: **; p < 0.001: ***; p < 0.0001: ****.

**Suppl. Fig. 6**. Dice and Jaccard coefficients, Hausdorff distance, and absolute volume difference of ICV segmentation on downsampled adversarial cases. not significant: ns, p < 0.05: *; p < 0.01: **; p < 0.001: ***; p < 0.0001: ****.

**Suppl. Fig. 7**. Dice and Jaccard coefficients, Hausdorff distance, and absolute volume difference of ventricular segmentation on adversarial cases with lower resolution. not significant: ns, p < 0.05: *; p < 0.01: **; p < 0.001: ***; p < 0.0001: ****.

**Suppl. Fig. 8**. Dice and Jaccard coefficients, Hausdorff distance, and absolute volume difference of ventricular segmentation on adversarial cases with increased noise. not significant: ns, p < 0.05: *; p < 0.01: **; p < 0.001: ***; p < 0.0001: ****.

