## Supplementary Material for "Improved segmentation of the intracranial and ventricular volumes in populations with cerebrovascular lesions and atrophy using 3D CNNs"

**Tables**

**Suppl. Table 1. Average Dice coefficient for brain and ventricular segmentation across models with different noise-induced sequences**

| **Region of Interest** | **Noisy Sequence** | **Dice Coefficient** |
| --- | --- | --- |
| Whole brain | T1 | **0.982 ± 0.015** |
|  | T2 | 0.983 ± 0.006 |
|  | FLAIR | 0.984 ± 0.003 |
| Ventricles | T1 | 0.892 ± 0.049 |
|  | T2 | 0.910 ± 0.041 |
|  | FLAIR | 0.910 ± 0.041 |

**Suppl. Table 2. Average Dice coefficient for brain and ventricular segmentation across models with different sequence variations**

| **Structure** | **Sequences** | **Validation Dice coefficient** |
| --- | --- | --- |
| Whole brain | T1+T2+FLAIR | **0.988** |
|  | T1+T2 | 0.982 |
|  | T1+FL | 0.985 |
|  | T1 | 0.987 |
| Ventricles | T1+T2+FLAIR | **0.910** |
|  | T1+T2 | 0.897 |
|  | T1+FLAIR | 0.836 |
|  | T1 | 0.947 |

**Suppl. Table 3. Dice coefficients for brain segmentation after post processing using a CRF**

| Smoothness Kernel | Appearance Kernel | Homogeneity | Dice Coefficient |
| --- | --- | --- | --- |
| 1, 1, 1, 1 | 3, 5, 5, 5 | 4.0, 4.0, 4.0 | **0.993 ± 0.009** |
| 3, 3, 3, 3 | 17, 12, 10, 5 | 4.5, 4.0, 3.0 | 0.948 ± 0.014 |
| 5, 5, 5, 5 | 17, 12, 10, 5 | 5.5, 5.0, 4.5 | 0.928 ± 0.018 |
| 1, 1, 1, 1 | 17, 12, 10, 5 | 4.0, 4.0, 4.0 | 0.947 ± 0.018 |
| 3, 3, 3, 3 | 3, 5, 5, 5 | 4.0, 4.0, 4.0 | 0.979 ± 0.006 |

**Suppl. Table 4. Dice coefficients for ventricle segmentation after post processing using a CRF**

| **Smoothness Kernel** | **Appearance Kernel** | **Homogeneity** | **Dice Coefficient** |
| --- | --- | --- | --- |
| 1, 1, 1, 1 | 3, 5, 5, 5 | 4.0, 4.0, 4.0 | **0.954 ± 0.019** |
| 3, 3, 3, 3 | 17, 12, 10, 5 | 4.5, 4.0, 3.0 | 0.922 ± 0.030 |
| 5, 5, 5, 5 | 17, 12, 10, 5 | 5.5, 5.0, 4.5 | 0.890 ± 0.051 |
| 1, 1, 1, 1 | 17, 12, 10, 5 | 4.0, 4.0, 4.0, 4.0 | 0.936 ± 0.020 |
| 3, 3, 3, 3 | 3, 5, 5, 5 | 4.0, 4.0, 4.0 | 0.948 ± 0.021 |

**Suppl. Table 5. Clinical adversarial attacks comprising downsampling and addition of multiplicative noise**

| **Structure** | **Input** | **Attack** | **Dice coefficient drop** |
| --- | --- | --- | --- |
| **Whole Brain** | Multi-contrast | Downsampling to 2 iso | < 0.1% |
|  |  | Downsampling to 2x2x4 | 0.2% |
|  |  | Noise σ=0.1 | 0.1% |
|  |  | Noise σ=0.3 | 0.2% |
|  |  | Noise σ=0.5 | 0.1% |
|  | T1-only | Downsampling to 2 iso | 4.6% |
|  |  | Downsampling to 2x2x4 | 4.7% |
|  |  | Noise σ=0.1 | 0.6% |
|  |  | Noise σ=0.3 | 0.7% |
|  |  | Noise σ=0.5 | 3% |
| **Ventricles** | Multi-contrast | Downsampling to 2 iso | 1.8% |
|  |  | Downsampling to 2x2x4 | 2.4% |
|  |  | Noise σ=0.1 | < 0.1% |
|  |  | Noise σ=0.3 | 1.5% |
|  |  | Noise σ=0.5 | 3.8%* |
|  | T1-only | Downsampling to 2 iso | 1.7% |
|  |  | Downsampling to 2x2x4 | 1.9% |
|  |  | Noise σ=0.1 | 0.2% |
|  |  | Noise σ=0.3 | 1.5% |
|  |  | Noise σ=0.5 | 3.1% |

**Suppl. Table 6.** Dice coefficients, Jaccard coefficients, Hausdorff distances, and absolute volume differences of ICV segmentation methods on adversarial cases with lower resolution

|  | **iCVMapper**  (Multi-  contrast) | **iCVMapper**  (T1w only) | **MONSTR**  (Multi-  contrast) | **BET**  (Multi-  contrast) | **FreeSurfer***  (T1w only) |
| --- | --- | --- | --- | --- | --- |
| **Dice coefficient** | 0.980 ± 0.006 | 0.866 ± 0.096 | 0.603 ± 0.117 | 0.744 ± 0.162 | 0.928 ± 0.019 |
| **Jaccard coefficient** | 0.960 ± 0.011 | 0.775 ± 0.132 | 0.441 ± 0.118 | 0.618 ± 0.192 | 0.866 ± 0.031 |
| **Hausdorff Distance (mm)** | 2.896 ± 3.522 | 26.391 ± 20.144 | 42.224 ± 13.001 | 44.469 ± 27.105 | 4.685 ± 1.300 |
| **Absolute Volume Difference (%)** | 7.746 ± 6.227 | 17.767 ± 18.622 | 3.840 ± 4.010 | 47.780 ± 54.566 | 2.954 ± 2.410 |

*FreeSurfer failed to process 7 scans.

**Suppl. Table 7.** Dice coefficients, Jaccard coefficients, Hausdorff distances, and absolute volume differences of ventricular segmentation methods on adversarial cases with lower resolution

|  | **VentMapper** (Multi-contrast) | **VentMapper**  (T1-based) | **FreeSurfer***  (T1-based) |
| --- | --- | --- | --- |
| **Dice coefficient** | 0.922 ± 0.038 | 0.927 ± 0.035 | 0.886 ± 0.044 |
| **Jaccard coefficient** | 0.858 ± 0.063 | 0.866 ± 0.058 | 0.798 ± 0.069 |
| **Hausdorff distance (mm)** | 2.896 ± 3.522 | 2.623 ± 1.169 | 4.685 ± 1.300 |
| **Absolute Volume Difference (%)** | 7.747 ± 6.227 | 7.659 ± 6.150 | 2.954 ± 2.410 |

*FreeSurfer failed to process 7 scans.

**Suppl. Table 8.** Dice coefficients, Jaccard coefficients, Hausdorff distances, and absolute volume differences of ICV segmentation methods on adversarial cases with increased noise

|  | **iCVMapper**  (Multi-  contrast) | **iCVMapper**  (T1w only) | **MONSTR***  (Multi-  contrast) | **BET**  (Multi-  contrast) | **FreeSurfer***  (T1w only) |
| --- | --- | --- | --- | --- | --- |
| **Dice coefficient** | 0.981 ± 0.014 | 0.958 ± 0.017 | 0.599 ± 0.113 | 0.177 ± 0.214 | 0.386 ± 0.149 |
| **Jaccard coefficient** | 0.963 ± 0.025 | 0.921 ± 0.030 | 0.437 ± 0.117 | 0.116 ± 0.166 | 0.251 ± 0.125 |
| **Hausdorff Distance (mm)** | 2.907 ± 4.321 | 6.044 ± 4.307 | 42.011 ± 13.037 | 69.010 ± 18.893 | 40.287 ± 15.821 |
| **Absolute Volume Difference (%)** | 2.235 ± 3.471 | 3.626 ± 3.551 | 3.690 ± 3.690 | 84.940 ± 21.693 | 68.463 ± 28.613 |

*FreeSurfer failed to process 42 scans and MONSTR failed to process 5 scans.

**Suppl. Table 9.** Dice coefficients, Jaccard coefficients, Hausdorff distances, and absolute volume differences of ventricular segmentation methods on adversarial cases with increased noise

|  | **VentMapper** (Multi-contrast) | **VentMapper**  (T1-based) | **FreeSurfer***  (T1-based) |
| --- | --- | --- | --- |
| **Dice coefficient** | 0.915 ± 0.033 | 0.909 ± 0.036 | 0.154 ± 0.247 |
| **Jaccard coefficient** | 0.847 ± 0.053 | 0.835 ± 0.057 | 0.110 ± 0.199 |
| **Hausdorff distance (mm)** | 1.884 ± 4.282 | 1.785 ± 0.759 | 40.287 ± 15.821 |
| **Absolute Volume Difference (%)** | 4.937 ± 4.867 | 6.402 ± 4.227 | 60.464 ± 28.614 |

*FreeSurfer failed to process 42 scans.

**Figures**

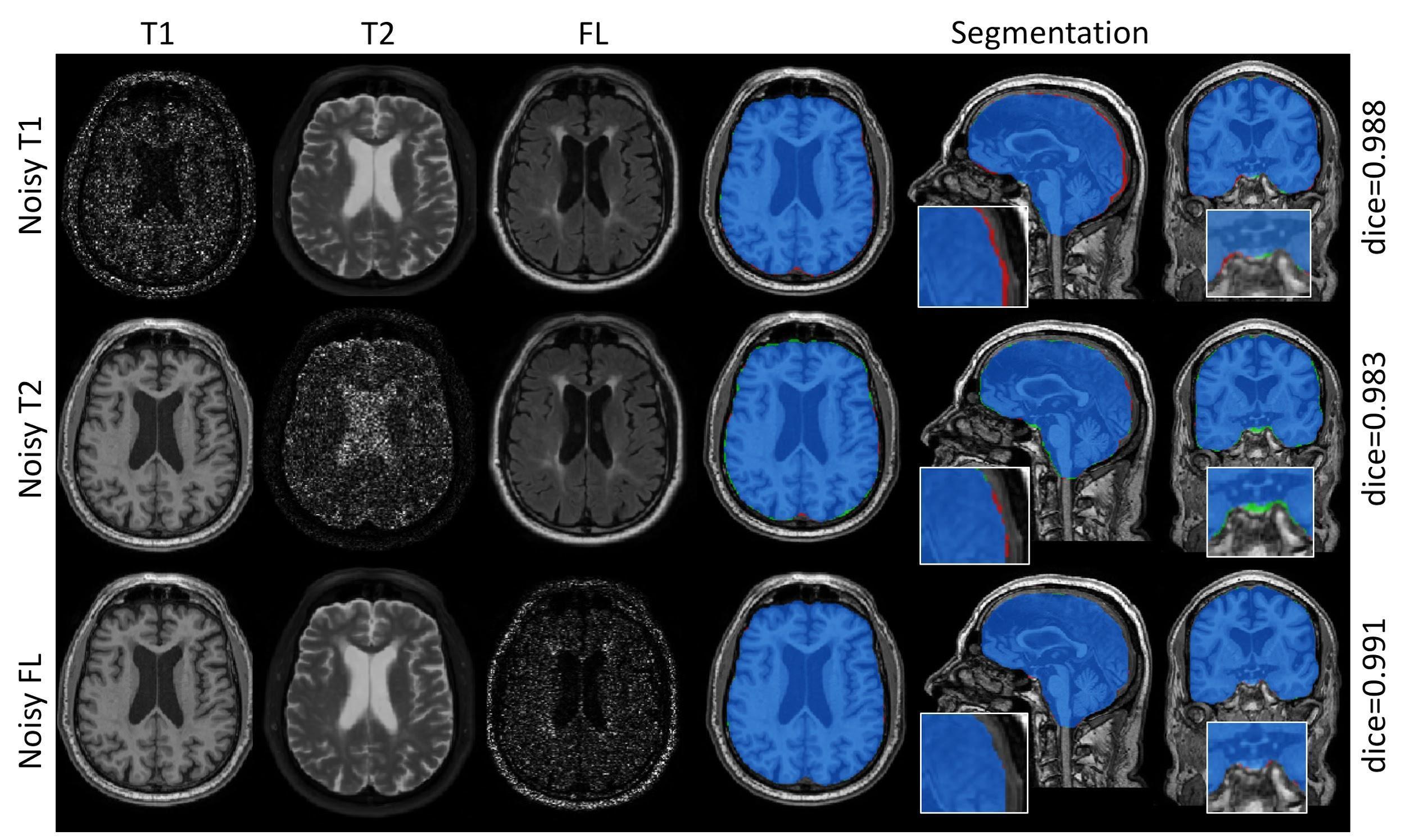

**Suppl. Fig. 1.** Effect of input sequence (contrast) on ICV segmentation accuracy. Blue represents overlap, Red voxels present in ground truth missing in prediction, Green voxels present in prediction and not in ground truth.

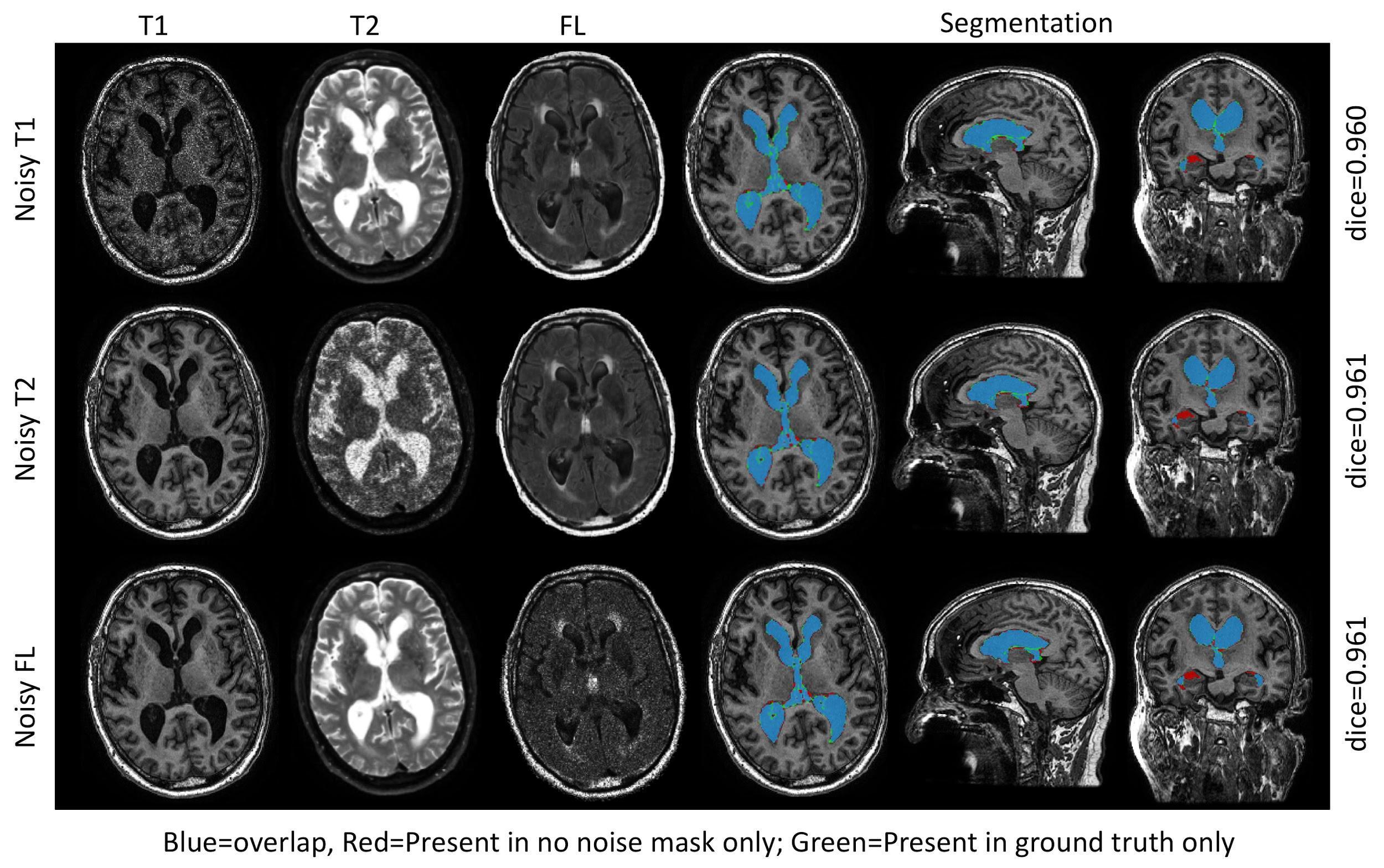

**Suppl. Fig. 2.** Effect of input sequence (contrast) on ventricular segmentation accuracy. Blue represent overlap, Red voxels present in ground truth missing in prediction, Green voxels present in prediction and not in ground truth.

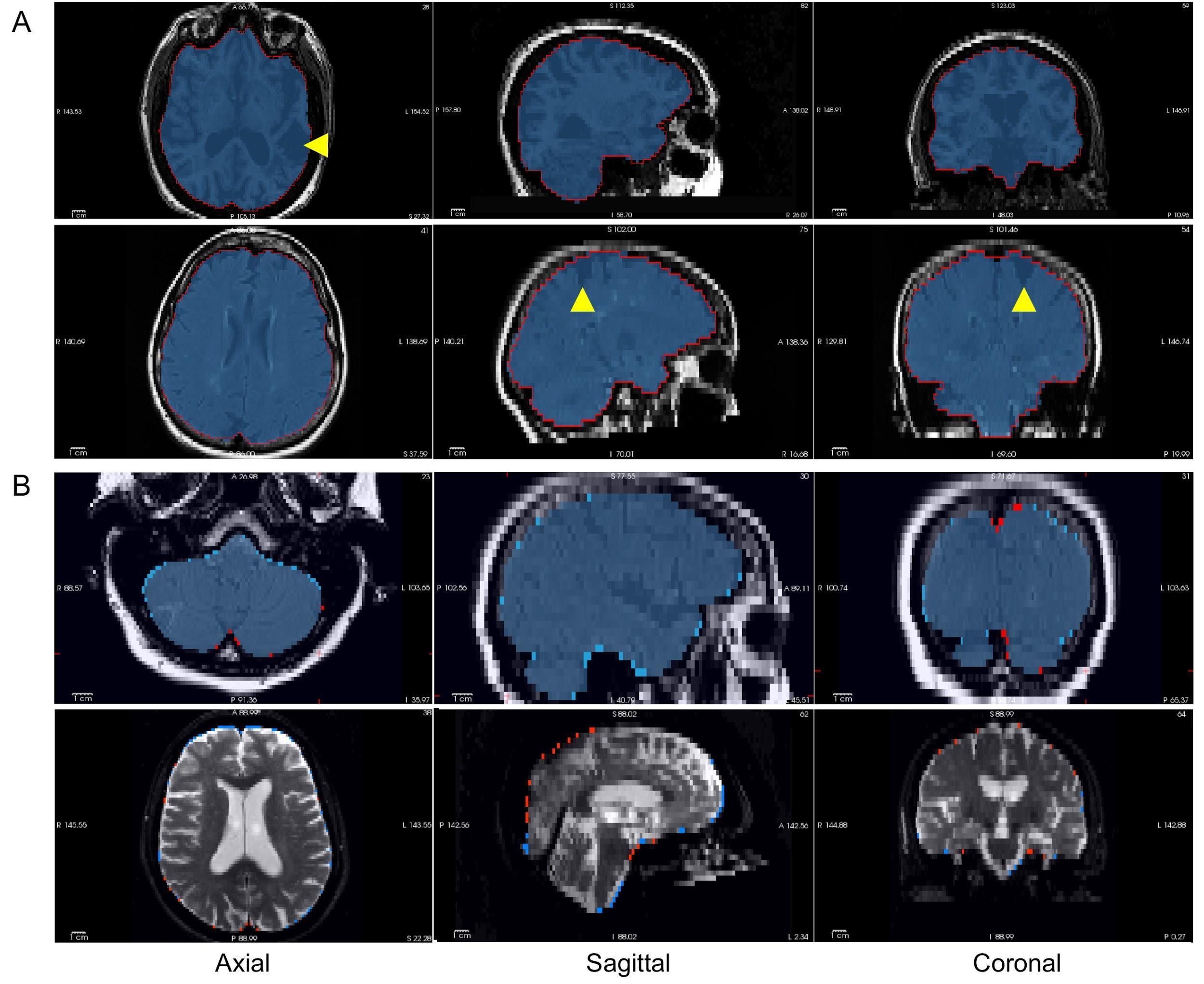

**Suppl. Fig. 3.** Brain segmentation in four test cases from different studies highlighting the quality of segmentations. Yellow arrowheads show strokes.

(A) Predicted segmentations (blue label) overlaid on manual delineations (red outline).

(B) Mis-segmented voxels highlight the differences between the segmentations. Red voxels are manually labeled voxels not predicted by the model, while light blue voxels are predicted voxels that were not present in the manual labels.

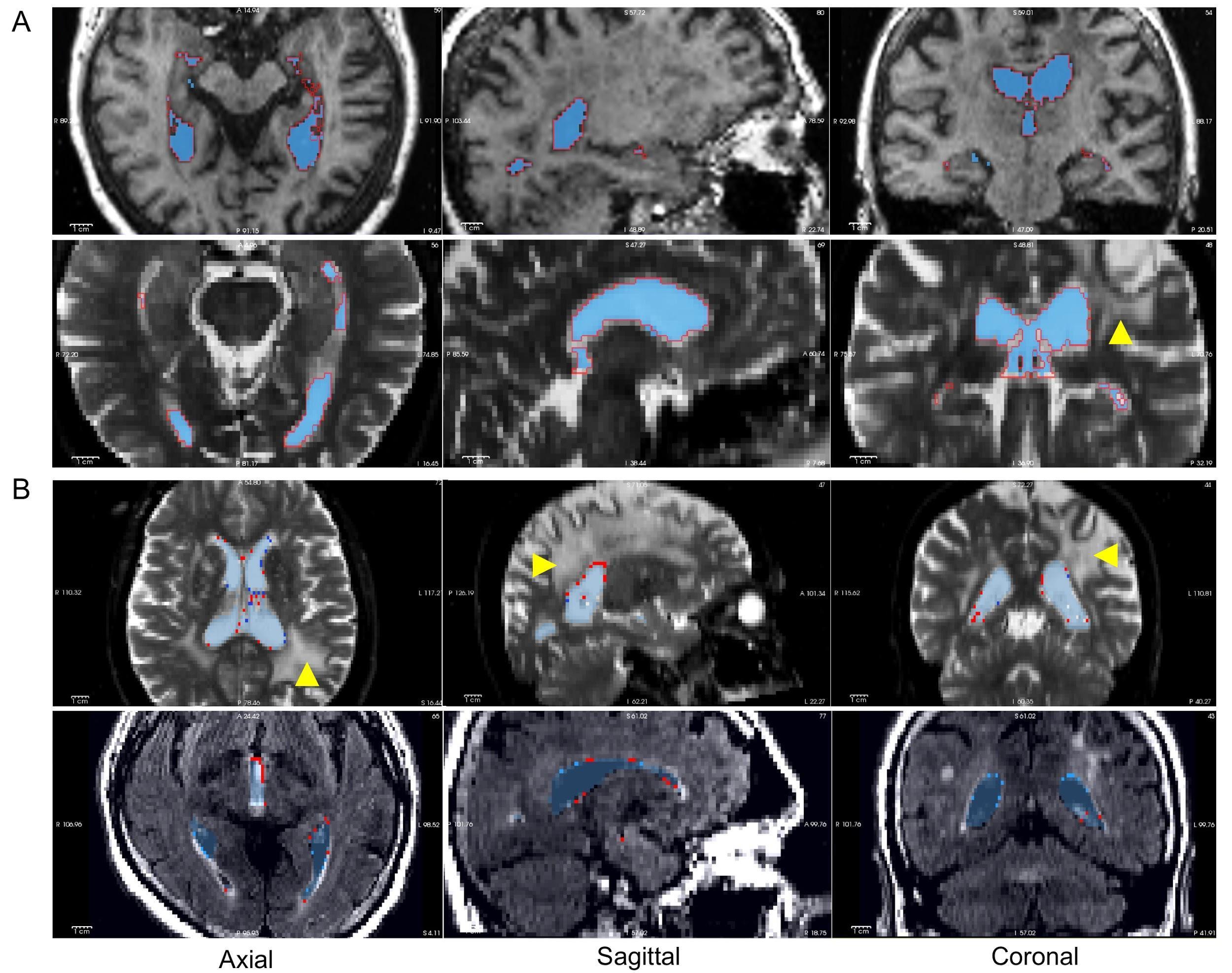

**Suppl. Fig. 4.** Ventricular segmentation in four test cases from different studies highlighting the quality of segmentations. Yellow arrowheads show severe white matter hyperintensities (WMH) or stroke.

(A) Shows under segmentation of our model (blue label) by overlay on manual delineations (red).

(B) Shows the overlap in our model compared to the manual segmentation by voxels incorrectly labelled by our model that were not in the manual segmentation (light blue) and voxel missed by our model that were present in the manual segmentation (red).

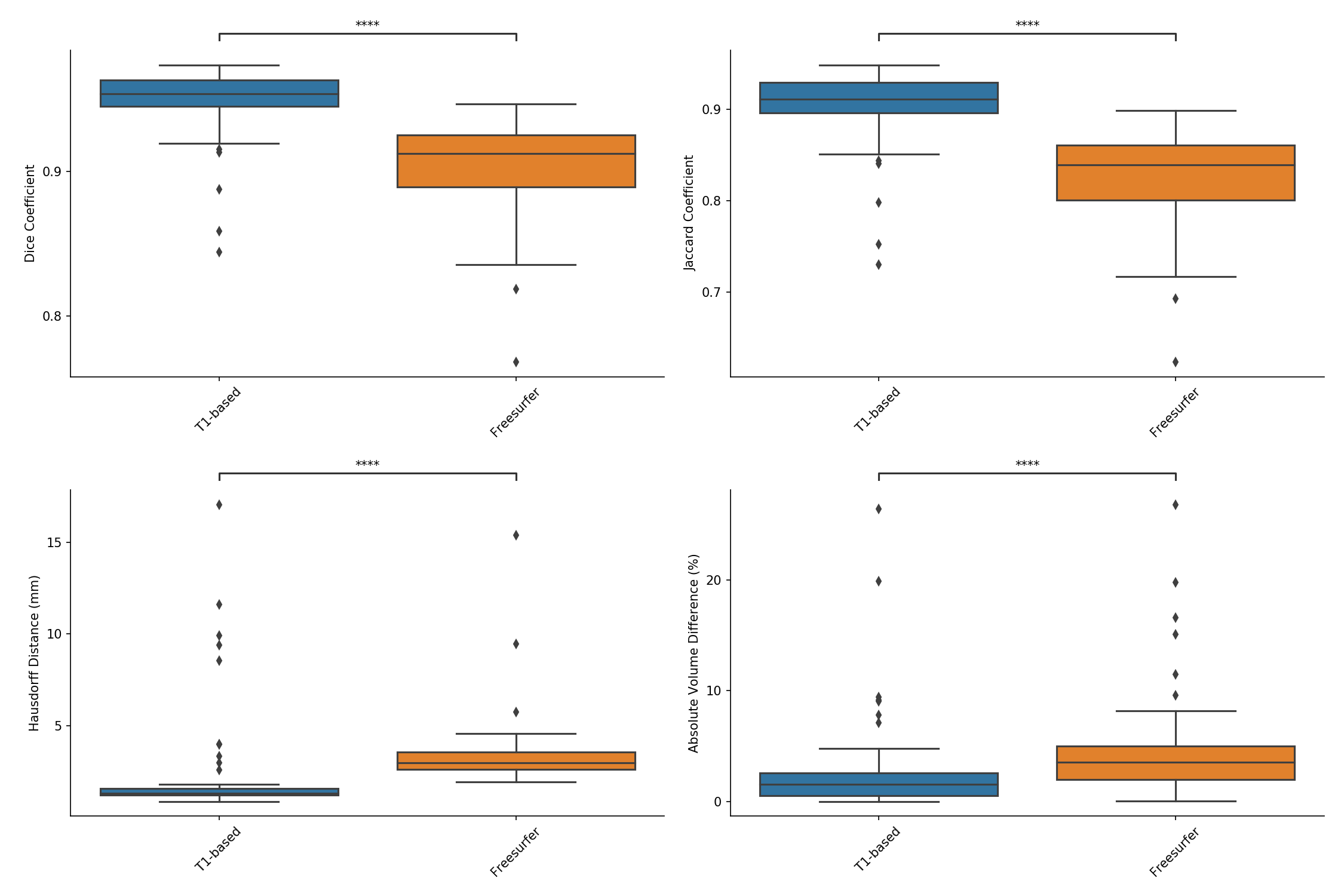

**Suppl. Fig. 5.** Dice and Jaccard coefficients, Hausdorff distance, and absolute volume difference of ventricular segmentation on the SDS vascular cognitive impairment cohort. not significant: ns, p < 0.05: *; p < 0.01: **; p < 0.001: ***; p < 0.0001: ****.

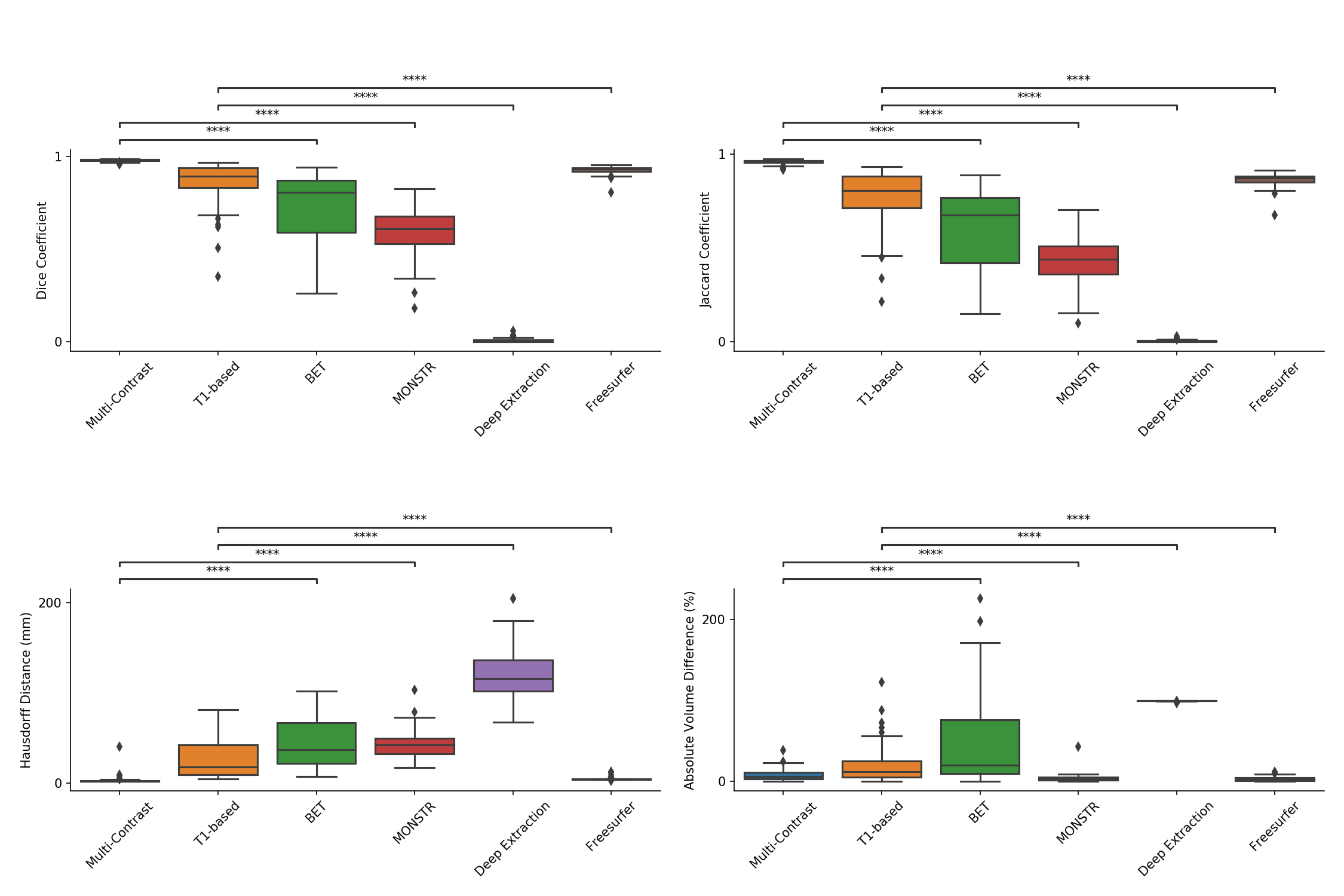

**Suppl. Fig. 6.** Dice and Jaccard coefficients, Hausdorff distance, and absolute volume difference of ICV segmentation on adversarial cases with lower resolution. not significant: ns, p < 0.05: *; p < 0.01: **; p < 0.001: ***; p < 0.0001: ****.

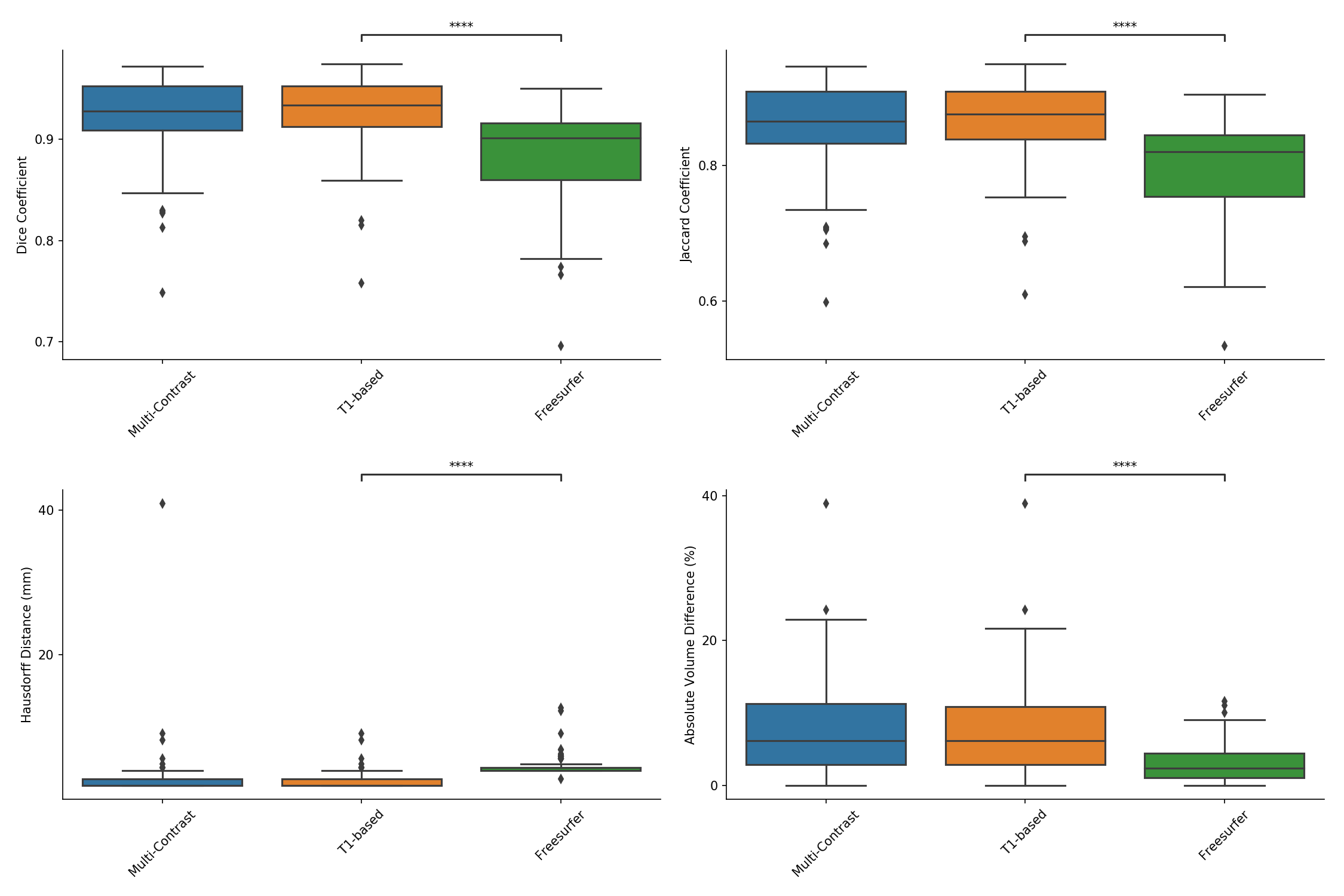

**Suppl. Fig. 7.** Dice and Jaccard coefficients, Hausdorff distance, and absolute volume difference of ventricular segmentation on adversarial cases with lower resolution. not significant: ns, p < 0.05: *; p < 0.01: **; p < 0.001: ***; p < 0.0001: ****.

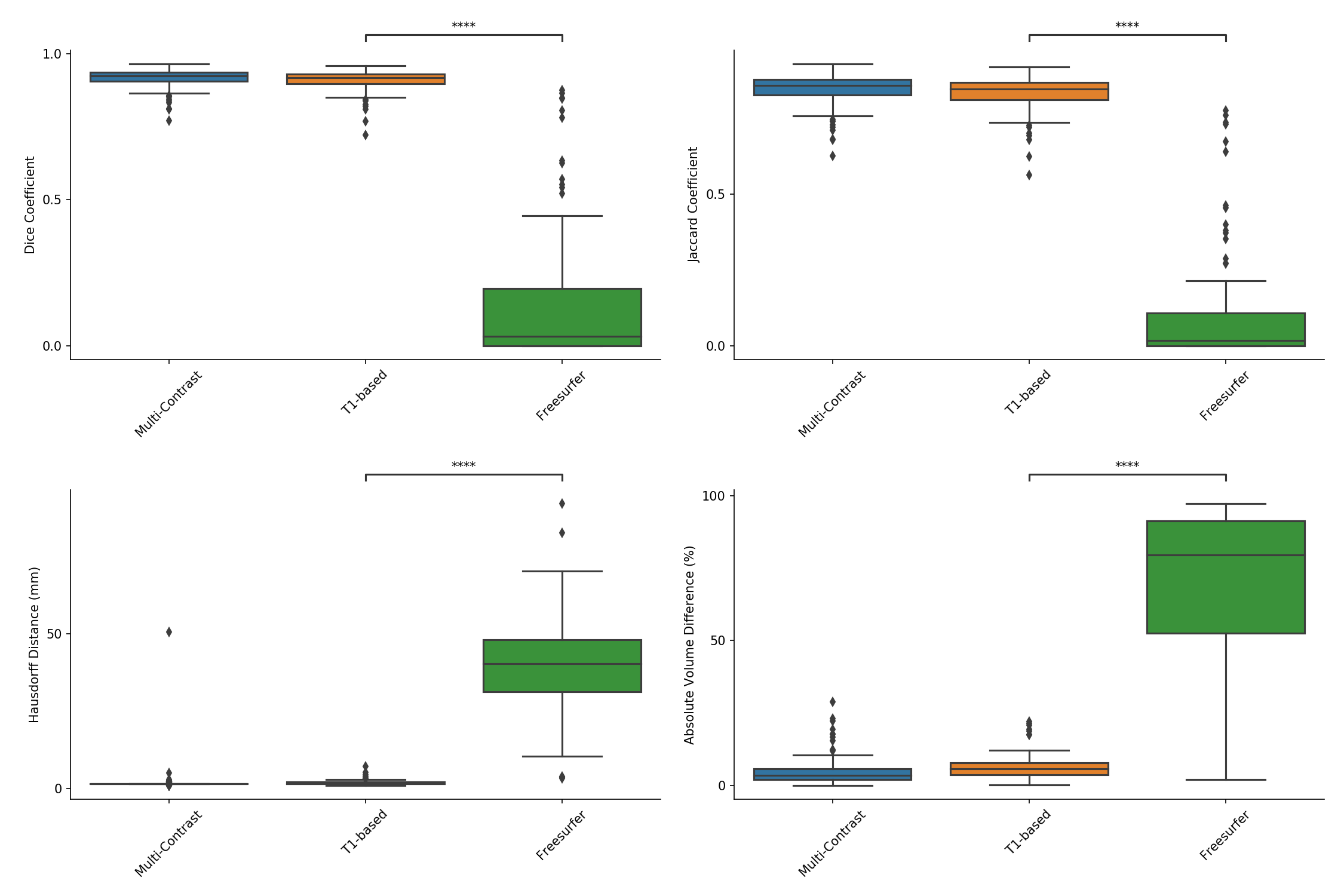

**Suppl. Fig. 8.** Dice and Jaccard coefficients, Hausdorff distance, and absolute volume difference of ventricular segmentation on adversarial cases with increased noise. not significant: ns, p < 0.05: *; p < 0.01: **; p < 0.001: ***; p < 0.0001: ****.

**MRI acquisition parameters**

*Ontario Neurodegenerative Disease Research Initiative (ONDRI),*

*Medical Imaging Trials NEtwork of Canada (MITNEC) Project C6:*

3DT1: TE = (GE: Min full, Philips: 2.3 ms, Siemens: 2.98 ms); TR = (GE: Min, Philips: 9.5 ms, Siemens: 2300 ms); TI = (GE: 400 ms, Philips: 925 ms, Siemens: 900 ms); slice thickness = 1 mm; matrix size = 256 x 256; in-plane field of view [FOV] = 256 x 256 mm.

T2: TE = (GE: Min full, Philips: 10 ms, Siemens: 10 ms), TR = 3000 ms, slice thickness = 3 mm, matrix size = (GE: 256 x 256, Philips: 256 x 234, Siemens: 256 x 256), in-plane FOV = 240 x 240 mm, phase FOV= (GE: 75%, Philips: 75%, Siemens: 81%).

2D FLAIR: TE = (GE: 140 ms, Philips: 90 ms, Siemens: 120 ms), TR = 9000, slice thickness = 3 mm, matrix = (GE: 256 x 256, Philips: 256 x 234, Siemens: 256 x 256), field of view [FOV] = 240 x 240.

*Canadian Atherosclerosis Imaging Network (CAIN):*

3DT1: TE = 2.3 ms; TR = 9.5 ms; TI = 1400 ms; slice thickness = 1.4 mm; matrix size = 256 x 164.

T2: TE = 10.7/102 ms; TR = 2500 ms; TI = 0 ms; slice thickness = 3 mm; matrix size = 256 x 216.

T2 FLAIR: TE = 2.3 ms; TR = 9000 ms; TI = 2800 ms; slice thickness = 3 mm; matrix size = 256 x 164.

*Sunnybrook Dementia Study (SDS):*

3DT1: TE: 5 ms; TR: 35 ms; TI: 35 ms; slice thickness = 1.2 mm; matrix size = 256 x 192.
